# Can the Z:A ratio serve as a genomic index of sexual selection? An assessment across space and time in a genus of lekking birds

**DOI:** 10.64898/2026.07.21.739449

**Authors:** H. Luke Anderson, Kevin F. P. Bennett, Kira M. Long, Jeffrey D. Brawn, Aaron O’Dea, Thomas J. Parsons, Philip L. F. Johnson, Michael J. Braun

## Abstract

Understanding the role of sexual selection in shaping evolutionary trajectories is a key goal of biology, and thus developing a genomic measure of sexual selection strength is desirable. In systems with ZW sex determination, the ratio of nucleotide diversity (*π*) on the male-biased Z chromosome to that on the autosomes (the Z:A ratio) is expected to decline as male effective population size decreases under strong sexual selection, offering one such measure. However, non-equilibrium processes other than sexual selection (e.g., hybridization, changes in population size) can influence levels of Z-linked diversity relative to the rest of the genome, complicating interpretation of this metric and necessitating a systematic evaluation of how Z:A ratios vary across known demographic and selective circumstances. We leveraged longitudinal sampling over a ∼25-year period in *Manacus* manakin populations in western Panama, spanning a mainland transect (including *M. candei*, *M. vitellinus*, and their hybrid zone) and insular populations of the Bocas del Toro Archipelago. On the mainland, Z:A ratios were temporally stable within populations, but varied across populations: (1) Z:A ratios were lower in *M. vitellinus* than *M. candei*, likely due to stronger sexual selection in the former; and (2) Z:A ratios increased at the hybrid zone center, likely due to fast-Z effects in parentals inflating Z:A ratios in recent hybrids. Island populations exhibited significantly lower Z:A ratios than mainland populations, consistent with theoretical expectations of Z:A reductions following population contractions as the islands became isolated. Insular Z:A ratios increased with proximity to other landmasses, suggesting gene flow mitigates these effects in this system. While the Z:A ratio behaved predictably in relation to known non-equilibrium processes, our results demonstrate the potentially strong effects of demographic history and admixture on Z-linked diversity, which should be considered in studies aiming to use the Z:A ratio as a measure of sexual selection.

## INTRODUCTION

Sexual selection is a potent evolutionary force arising from competition over limited reproductive opportunities. Given the major role sexual selection can play in molding phenotypes and influencing speciation in nature (Andersson, 1994; Darwin, 1871), biologists are often interested in quantifying its contribution to the evolutionary process (e.g., Janicke et al., 2018). Common approaches for estimating sexual selection strength include the use of morphological or behavioral proxies (e.g., sexual dichromatism, sexual size dimorphism, display or vocalization complexity, mating system; Barber et al., 2024; Cally et al., 2021; Kraaijeveld et al., 2011; Nadeau et al., 2007; Seddon et al., 2013), as well as direct estimates of reproductive skew through observation of mating success or genetic analysis of paternity (e.g., Mackenzie et al., 1995; Ryder et al., 2009), which can then be used to calculate non-trait-based metrics of sexual selection (reviewed in Anthes et al., 2017). However, given the potentially limited correlation between phenotypic proxies and male-biased sexual selection *per se* (Burns, 1998; Dale et al., 2015; Irwin, 1994; Shultz & Burns, 2017), the difficulty of obtaining field-based estimates of reproductive skew for large numbers of species, and the fact that short-term field studies are snapshots that may not be representative of selection over evolutionary timescales, developing alternative methods to assess the strength of sexual selection in wild populations is desirable.

Recent advances in genomic technology have enabled powerful inferences about the evolutionary forces shaping natural populations, including sexual selection. One promising index of sex-biased reproductive skew can be obtained by estimating the effective population size (*N_e_*) of sex-linked genomic compartments relative to the autosomes. In organisms with ZW chromosomal sex determination (e.g., birds, lepidopterans, some reptiles and fish), where males are homogametic (ZZ) and females are heterogametic (ZW), the *N_e_* of the male-biased Z chromosome is expected to be ¾ that of the autosomes. This is because, in a population with equal sex ratios, there are three copies of a Z locus for every four copies of an autosomal locus. Given that the nucleotide diversity (π) of each chromosome is directly proportional to its *N_e_*, the ratio of nucleotide diversity on the male-biased Z chromosome relative to the autosomes (hereafter, the Z:A ratio) is expected to be 0.75 under neutrality (Charlesworth, 2009; Corl & Ellegren, 2012; Irwin, 2018). Male-biased sexual selection reduces the *N_e_* of males relative to females, leading to disproportionate reductions of *N_e_*Z relative to *N_e_*A, and reductions of the Z:A ratio below the neutral expectation (Figure 1; Charlesworth, 2001, 2009). Thus, the Z:A ratio should offer a measure of the degree of male reproductive skew in a population.

**Figure 1.**
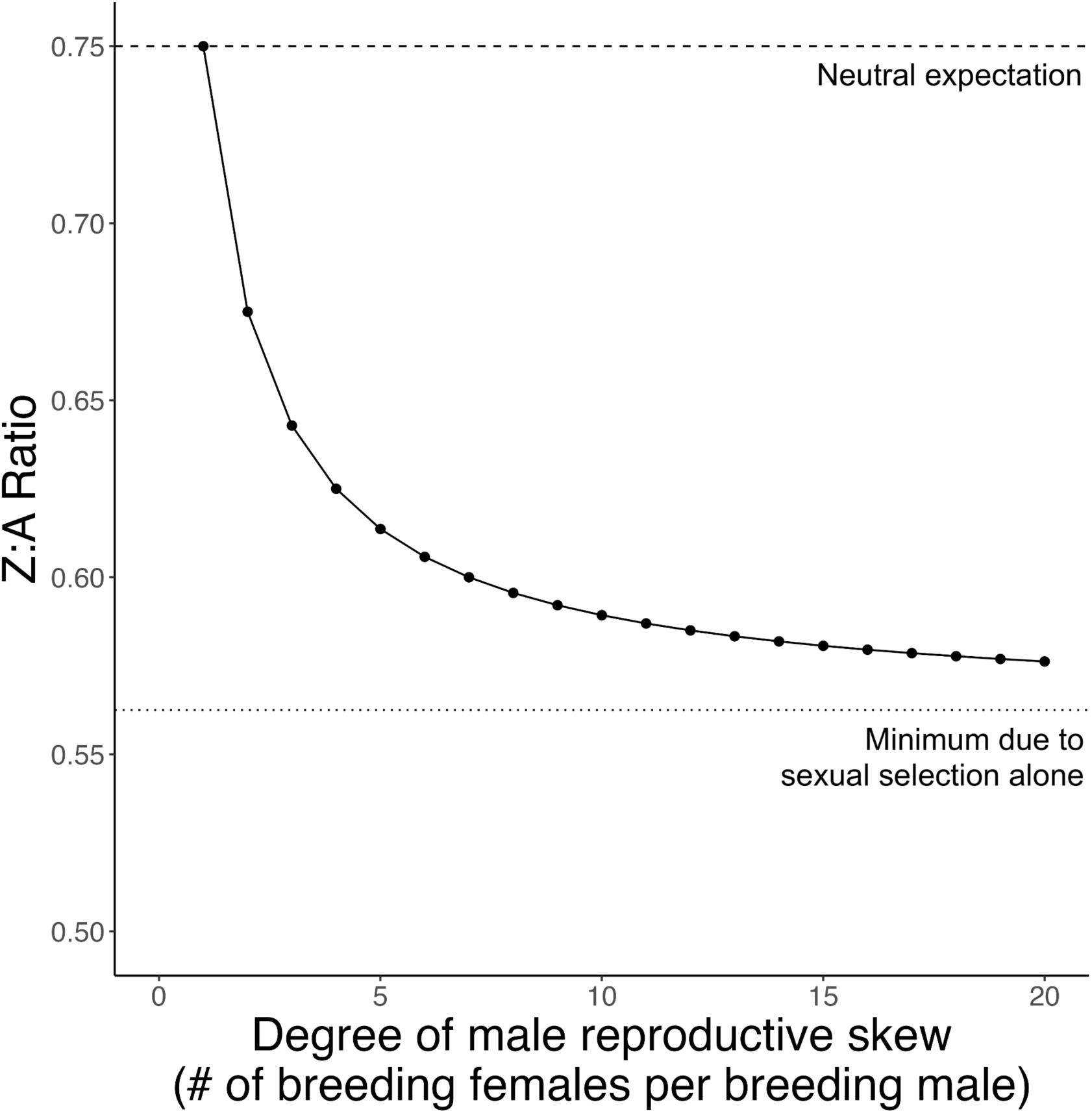
Theoretical relationship between the strength of male-biased sexual selection and the Z:A ratio. The nucleotide diversity (π) of a given genomic compartment is expected to be directly proportional to the effective population size (*N_e_*) of that compartment. As such, the Z chromosome, which occurs twice in males but only once in females, is expected to have ¾ the nucleotide diversity of the average autosome under equilibrium conditions (dashed line). As the degree of male reproductive skew increases, the effective population size of the Z chromosome (*N_e_Z*) relative to that of the autosomes (*N_e_A*) decreases, thereby reducing the Z_*π*_:A_*π*_ ratio. Sexual selection alone under the most extreme polygyny can only reduce *N_e_Z:N_e_A* to 9/16 (0.5625), per the equation *N_e_Z:N_e_A* = [(9*N*m*N*f)/(2*N*m +4*N*f)] / [(4*N*m*N*f)/(*N*m + *N*f)], where *N_m_* and *N_f_* represent the number of breeding males and females, respectively (Charlesworth 2001; Corl & Ellegren, 2012). Thus, the curve asymptotes to 0.5625 (dotted line). Reductions below this value require the influence of additional factors (e.g., demographic or selective events, life history differences between males and females, etc.).

Despite these relatively straightforward theoretical expectations, empirical data paint a more complex picture. Studies in birds—a clade replete with elaborate male sexual ornaments and behaviors, as well as a rich history of research in sexual selection—have yielded valuable but sometimes ambiguous information about the degree to which the Z:A ratio may serve as a reliable index of sexual selection strength. For example, a study examining phylogenetic pairs of shorebirds found that polygynous species had lower Z:A ratios than monogamous species, consistent with the expectation of greater male reproductive skew under polygyny (Corl & Ellegren, 2012). Similarly, a recent study found that polygynous manakins exhibited lower Z:A ratios than two socially monogamous outgroup taxa (although differences in Z:A ratios among manakin species did not always align with expectations; Balakrishnan et al., 2026). However, a study examining monochromatic and dichromatic species pairs across a range of avian families yielded a surprising result: monochromatic species had consistently lower Z:A ratios than dichromatic ones (Huang & Rabosky, 2015), contrary to long-held expectations that plumage dichromatism should serve as an indicator of male-biased sexual selection (Andersson, 1994; Darwin, 1871). Given these results, and the degree to which methodological differences across studies can complicate interpretation and generalization (Irwin, 2018), the factors shaping sex chromosome diversity require more systematic scrutiny before the Z:A ratio can be confidently applied as an index of sexual selection strength.

Of note, sex-specific mutational, life history, and selective factors aside from male reproductive skew can influence the genetic diversity of the Z chromosome. Some factors (e.g., male-biased mutation, longer male generation times) are generally expected to increase Z-linked diversity relative to the autosomes, whereas others are expected to deplete it (e.g., efficient selection in hemizygous state, reduced recombination). The Z chromosome is expected to have higher mutation rates than the autosomes because it occurs twice as frequently in males as in females, and males experience higher germline mutation rates than females due to the frequency of meiotic divisions associated with sperm production (Wilson Sayres, 2018; Wilson Sayres et al., 2011; Wilson Sayres & Makova, 2011; Axelsson et al., 2004; Wang et al., 2014). Differences in male and female generation times—common in birds, where males may exhibit delayed plumage maturation and/or reproductive success (e.g., Schaedler et al., 2021)— can also influence sex-linked diversity, as species with higher male:female generation time ratios will accumulate relatively more mutations on the male-biased sex chromosome over evolutionary timescales (Amster & Sella, 2016, 2020). In contrast, more efficient positive and purifying selection on the Z chromosome (due to the exposure of recessive alleles to selection in the heterogametic sex) should deplete Z-linked genetic diversity, effects that may be amplified by genetic hitchhiking and background selection, respectively. Moreover, because the Z chromosome only recombines in males, lower recombination rates on Z mean that large hitchhiking blocks can persist longer (Cutter, 2019).

Demographic processes and events are also expected to have strong effects on sex-linked diversity. For example, sex-biased mortality, dispersal, or migration may all influence relative levels of sex-linked genetic diversity (Ellegren, 2009a), as can changes in census population size. Notably, coalescent modeling demonstrates that population size contractions should result in transient reductions in sex-linked diversity relative to the autosomes, as sex chromosomes reach new equilibrium values more rapidly than the rest of the genome due to their smaller *N_e_* (Pool & Nielsen, 2007). Indeed, the Z:A ratios reported for many passerine birds (e.g., Borge et al., 2005; Huynh et al., 2010; Janoušek et al., 2019; Oyler-McCance et al., 2015; Schield et al., 2021; Sundström et al., 2004; Irwin, 2018) are well below the theoretical minimum possible by sexual selection alone (0.5625; Charlesworth, 2001, 2009; Corl & Ellegren, 2012). Authors have often invoked historical changes in population size or selective sweeps to explain the observed values, yet clear empirical demonstrations of either mechanism are lacking (but see Arbiza et al., 2014 and Wall et al., 2002 for X:A examples in humans and *Drosophila*).

Admixture and hybridization may also affect Z:A ratios. The Z chromosome is known to evolve more rapidly than the autosomes (the “fast-Z effect”; Mank et al., 2007; Ellegren, 2009b) due to its higher mutation rate, smaller *N_e_* (potentially resulting in higher fixation of mildly deleterious variants; Mank et al., 2010; Wright et al., 2015; Wanders et al., 2024) and/or hemizygosity in females (increasing the efficacy of positive selection on recessive mutations; Charlesworth et al., 1987). As such, allopatric lineages are expected to evolve fixed differences on the Z more rapidly than on the autosomes. This increased divergence between parentals on the Z chromosome would result in elevated Z-linked heterozygosity (and higher Z:A ratios) in early-generation hybrids. Additionally, Z:A ratios in hybrid zones may be influenced by sex-biased mortality. Per Haldane’s Rule, hybrid inviability tends to manifest in the heterogametic sex (Haldane, 1922). Thus, in ZW systems, increased mortality in heterogametic females would be expected to reduce female *N_e_* and elevate Z:A ratios in hybrid populations.

Due to the complexity of the factors potentially influencing Z-linked genetic diversity, we aimed to assess the stability of the Z:A ratio across populations within a single closely related group, where multifarious phylogenetic and life history variables are largely controlled. We leveraged a well-studied species complex of manakins in western Panama, where data were available for 20 populations spanning a range of known selective and biogeographic situations, offering a promising opportunity to disentangle the influence of various factors on Z:A ratios. Manakins (Aves: Pipridae) are a charismatic clade of neotropical birds known for the acrobatic courtship displays and flashy plumage of males. Most species have a lek-based polygynous mating system in which a small fraction of males obtains reproductive success; as such, manakins are thought to be evolving under strong male-biased sexual selection (Prum, 1997). The two lekking manakin species considered here, *Manacus vitellinus* and *M. candei*, form a stable hybrid zone in western Panama (Long et al., 2024), in which there is evidence of reduced hatching success in hybrids (Long et al., 2025). While courtship displays are similar between the two species and among the most complex in manakins (Lindsay et al., 2015), there is evidence that sexual selection may be stronger in *vitellinus* than *candei*. Specifically, *vitellinus* males exhibit higher levels of aggression (McDonald et al., 2001), a more physiologically demanding display repertoire (Miles et al., 2018), and an additional plumage ornament likely favored by females (Stein & Uy, 2006b). This ornament—yellow collar color—has introgressed from *vitellinus* into some populations of *candei*, apparently via sexual selection (Parsons et al., 1993; Brumfield et al., 2001; Bennett et al., 2021; Lim et al., 2024). In addition to the prior introgression of yellow collar color, several *candei* populations experienced introgression of olive belly color, another putatively sexually selected male plumage trait of *vitellinus,* between survey timepoints (Long et al., 2024), raising the possibility that this recent wave of introgression was accompanied by shifts in male reproductive skew.

We first examined a set of 11 populations along a mainland transect comprising parental and hybrid populations of *Manacus candei* and *M. vitellinus*, most of which were sampled at two timepoints ∼25 years apart. Broadly, we hypothesized that Z:A ratios among mainland *Manacus* populations would be generally consistent within species, but influenced by admixture, sexual selection, and introgression dynamics. First, we hypothesized that populations near the genomic center of the *candei*–*vitellinus* hybrid zone would exhibit higher Z:A ratios, for the reasons mentioned above. Second, given that *vitellinus and candei* differ in sexually selected traits, we hypothesized that Z:A ratios would be lower in *vitellinus* compared to *candei*, consistent with stronger sexual selection in the former. Third, we expected to observe temporal perturbations in Z:A ratios in populations that experienced changes in sexual selection dynamics between historical and contemporary survey timepoints.

We then compared the nucleotide diversity and Z:A ratios of mainland populations to a set of nine insular populations on the Bocas del Toro Archipelago, a series of continental islands that were gradually isolated from mainland Panama over the past 10,000 years (O’Dea et al., 2026). Some islands are quite small (range 22–6,000 ha), and the census size of their *Manacus* populations has been dramatically reduced compared to the mainland. Thus, we hypothesized that island populations would exhibit reduced Z:A ratios relative to mainland populations. This is because the Z chromosome is expected to more rapidly reach a lower equilibrium genetic diversity value after a population contraction than the autosomes due to its smaller *N_e_* (Pool & Nielsen, 2007). Given that theoretical models indicate Z:A ratios should depend upon both the size and recency of population contraction (Pool & Nielsen, 2007), we also hypothesized that variables related to island size and age would jointly influence Z:A ratios. Finally, we hypothesized that an island’s spatial proximity to other landmasses would positively correlate with Z:A ratios due to the greater potential for gene flow from other populations, which are expected to harbor more diverse or divergent Z chromosomes relative to the islands.

Ultimately, by assessing Z:A ratios across space and time in a well-characterized system, we aim to provide a test of the utility of the Z:A ratio as an indicator of sexual selection strength, as well as the degree to which this metric is influenced by known demographic processes.

## MATERIALS AND METHODS

### Study system and sampling design

We obtained blood or tissue samples from 434 *Manacus* manakins (see Supplementary Materials for details) belonging to populations in two major geographic zones: (1) a mainland transect spanning parental populations of the white-collared manakin (*M. candei*), golden-collared manakin (*M. vitellinus*), and their hybrid zone in western Panama (hereafter, the “mainland transect” dataset); and (2) populations residing on the islands of the Bocas del Toro Archipelago (hereafter, the “islands” dataset; Figure 2).

**Figure 2.**
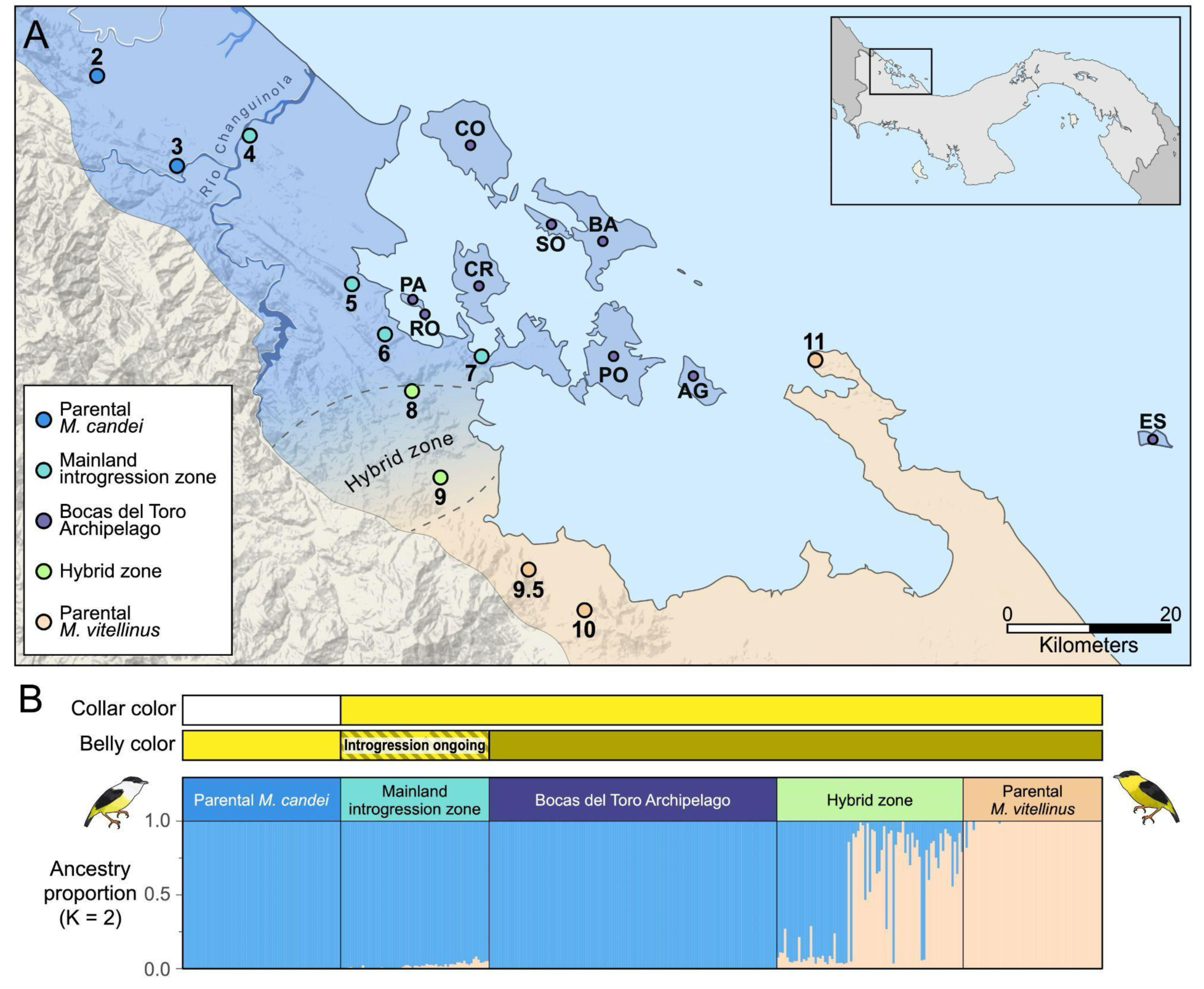
Map and ancestry of *Manacus* populations on mainland Panama and the Bocas del Toro Archipelago. **(A)** The genomic features of the hybrid zone between *M. vitellinus* and *M. candei* are represented with color shading on the map. Circular points represent sampling localities along the mainland transect and in the Bocas del Toro islands. The peach-colored area represents the range where birds are genomically *M. vitellinus*, the blue-colored areas represent the range where birds are genomically *M. candei*, and the peach–blue gradient between populations 8 and 9 approximates the narrow genomic cline where local populations transition from being genomically *vitellinus*-like to *candei*-like. The approximate bounds of the transition zone are denoted by the dashed lines encompassing populations 8 and 9. The region between this hybrid zone and the Río Changuinola to the northwest is an introgression zone, where birds exhibit yellow collars (due to an introgressed allele from *vitellinus*) but are predominantly genetically similar to *candei*. All mainland localities other than populations 7, 9.5, and 11 were sampled at both historical and contemporary timepoints. Islands sampled at least once include: Cayo de Agua (AG), Isla Bastimentos (BA), Isla Cristóbal (CR), Isla Colón (CO), Escudo de Veraguas (ES), Isla Pastores (PA), Cayo Roldan (RO), Isla Popa (PO), and Isla Solarte (SO); sampling localities on islands are not exact. Inset: Map of Panama, with the bounds of the main image indicated by the black box. **(B)** Horizontal bars represent the male collar and belly plumage phenotypes of the individuals across the different populations, and the parental phenotypes are illustrated by the bird cartoons. Belly color introgression was incomplete during historical sampling of the mainland introgression zone populations and has been ongoing (Long et al 2024). The admixture plot below represents the proportion of ancestry at *K* = 2 for all sampled individuals. Parental *M. candei* ancestry is represented in blue, while parental *M. vitellinus* ancestry is shown in peach. Individuals from the Bocas del Toro Archipelago are genomically similar to parental *candei*. Mainland introgression zone populations are predominantly *candei* with a small amount of ancestry from *vitellinus*; these birds exhibit the introgressed yellow collar (and typically olive belly) color derived from *M. vitellinus.* Individuals sampled in the hybrid zone show a variable but high degree of admixture between the two parental lineages.

The mainland transect comprised 11 populations, which we treat in four groups, from west to east, as follows: parental *M. candei* (populations 2 and 3); an introgression zone, where male plumage traits from *M. vitellinus* have introgressed into the *M. candei* genomic background (populations 4–7); a narrow genomic transition or hybrid zone, where allele frequencies for numerous genetic markers transition rapidly from *vitellinus*-like to *candei*-like and many individuals are highly admixed (populations 8 and 9); and parental *M. vitellinus* (populations 9.5, 10, and 11). Eight populations (all but populations 7, 9.5, and 11) were sampled at two timepoints nearly three decades apart, enabling us to assess the temporal stability of Z:A ratios within populations. Historical sampling was conducted between 1989 and 1994 (Brumfield et al. 2001), while contemporary sampling was conducted between 2017 and 2020 (Long et al. 2024). Population identification numbers are consistent with those found in Brumfield et al. (2001), Yuri et al. (2009), Parchman et al. (2013), Lim et al. (2024), and Long et al. (2024). Previous analyses of a RAD-seq dataset (Long et al., 2024) demonstrated that the location of the genomic center of the hybrid zone near population 9 has remained stable in the years between the historical and contemporary surveys, although the cline for olive belly plumage (another putatively sexually selected ornament derived from *vitellinus*) has shifted significantly westward through the introgression zone between survey timepoints.

To investigate the effect of population contractions on the Z:A ratio, we also analyzed genomic data from *Manacus* populations on islands of the Bocas del Toro Archipelago (Figure 2), a group of islands off the coast of western Panama that were gradually isolated from the mainland by sea level rise beginning ∼10.6 ka (Lambeck et al., 2014; O’Dea et al., 2026). Samples were collected in the years 1987–1991, 2007, and 2022 from the following islands: Cayo de Agua (AG), Isla Bastimentos (BA), Isla Colón (CO), Isla Cristóbal (CR), Isla Escudo de Veraguas (ES), Cayo García, Isla Pastores (PA), Isla Popa (PO), Cayo Roldan (RO), and Isla Solarte (SO). The lone individual sampled from the tiny islet of Cayo García was combined with the Isla Pastores population for analysis, as the former is too small to harbor a substantial resident manakin population and the two islands are connected by mangroves. Island populations were lumped across years of collection and only considered in spatial analyses.

To maximize the number of Z chromosomes sampled and simplify downstream bioinformatic processing, we restricted our analyses to males. Individuals that appeared in previous studies were sexed in the field based on definitive male plumage or, in the case of green-plumaged individuals, using the molecular sexing protocol outlined in Long et al. (2024). Newly sequenced samples were sexed using the program *SATC* (Nursyifa et al., 2022), which performs sex assignment by identifying sex-linked loci based on sequencing coverage.

### RAD-seq library preparation, genotyping, and admixture analysis

Following a modified version of a published laboratory protocol (Etter et al., 2011), we used restriction site-associated DNA sequencing (RAD-seq) to obtain reduced-representation genomic data for individuals from the mainland transect and island populations (see Supplementary Materials). Raw reads were demultiplexed using process_radtags in *Stacks* v. 2.66 (Rochette et al., 2019) and aligned to the *M. vitellinus* reference genome using *bwa* mem v. 0.7.17 (Li, 2013; Li & Durbin, 2009). We used the gstacks and populations modules in *Stacks* to assess data quality and filter iteratively; ultimately, sites needed to have > 6x mean coverage, be present in all populations, and be present in ≥ 80% of the individuals from a population. Full details of library preparation, sequencing, and data filtering are outlined in the Supplementary Materials, and the number of Z-linked, autosomal, and macrosomal loci retained for each population after filtering is summarized in Table S1.

We used admixture analysis of this extensive SNP dataset to graphically display the genomic affinities of all 20 island and mainland populations. Prior to running ADMIXTURE v. 1.3 (Alexander et al., 2009) at *K* = 2, the variant dataset was filtered using bcftools v. 1.19 to include only biallelic sites (Danecek et al., 2021) and pruned to reduce linkage disequilibrium using plink v. 1.90b7.2 (--indep-pairwise 50 5 0.2; Chang et al., 2015). Although hierarchical structure at *K* = 3 does exist within the mainland transect (Long et al. 2024), we focus here on *K* = 2 clusters to emphasize affinities with the parental species *candei* and *vitellinus*, whose genomic differentiation dominates genetic structure across this system. Significant hierarchical structure also exists within the island populations, which is the subject of a forthcoming manuscript.

### Statistical analyses

We calculated nucleotide diversity (*π*; Nei & Li, 1979) from all-sites VCF files using the software *pixy* (v. 2.0.0.beta14), which generates unbiased estimates of *π* while accounting for missing data (Korunes & Samuk, 2021). Scaffolds were assigned to chromosomes based on a previous analysis of synteny with the chromosome-level *Chiroxiphia lanceolata* reference assembly (NCBI RefSeq Assembly: GCF_009829145.1) (Lim et al. 2024). We subsequently renamed chromosomes according to the zebra finch (*Taeniopygia guttata*) reference genome (NCBI RefSeq Assembly: GCF_000151805.1). We calculated *π* across Z-linked and autosomal scaffolds and divided those values to obtain Z:A ratios.

We first tested whether Z:A ratio estimates were influenced by the autosome type used in the denominator, as would be expected given that microsomes generally have higher rates of recombination than macrosomes. Macrosomal scaffolds were defined as those that mapped to autosomes ≥ 20 Mb in length (per Damas et al., 2018) in *Chiroxiphia lanceolata*, with chromosomes numbered according to their mapping to zebra finch (Figure S1). In *Chiroxiphia*, the 14 macrosomes range in size from 156.3 to 20.7 Mb, while the 19 microsomes range from 19.3 to 0.7 Mb. The Z chromosome (75.4 Mb) is comparable to an intermediate-sized macrosome. In our data, the Z falls at roughly the median of the macrosomal distribution in terms of total sites sequenced (Figure S1). Consistent with expectations, Z:A ratios calculated using only macrosomes in the denominator were higher than ratios calculated using only microsomes, and ratios using all autosomes were intermediate (Figure S2). Due to these significant differences according to the autosome type used in calculations (ANOVA: F_2,54_ = 263.1, p < 0.0001), and because the Z chromosome likely exhibits more similar patterns of recombination and GC-biased gene conversion to similarly sized macrosomes (Wanders et al., 2024), we report Z:A values based on macrosomes for the remainder of our results. We note that this choice influences only the absolute values of Z:A ratio and π estimates; results and conclusions about factors affecting Z:A ratio variation among populations are similar regardless of the autosome type used in calculations.

We then assessed temporal (within-population) and spatial (between-population) variation in the Z:A ratio along the mainland transect. We calculated population-level Z:A ratios by dividing the Z-linked *π* by the macrosomal *π* for each population at each sampling timepoint, yielding point estimates of Z:A ratios. To obtain estimates of measurement uncertainty for statistical comparisons along the mainland transect, we calculated individual-level Z:A ratios (i.e., by calculating the Z-linked *π* and the autosomal *π* for each individual separately; akin to heterozygosity) for each mainland population and timepoint (see Supplementary Materials for justification). We also calculated individual-level Z:A ratios for island populations, although these were not used in statistical analyses.

All additional statistical analyses were conducted in RStudio v. 2026.01.1+403 (Posit Studios, 2026). To assess spatial variation in genomic variables across mainland transect populations of *Manacus*, we fit generalized linear mixed effects models (GLMMs) in *glmmTMB* (Brooks et al., 2017) with Student’s *t* error distributions to handle heavy tails. These models included Z:A ratios, macrosomal π, or Z-linked π as response variables; population (locality along the transect) as a fixed effect; and a unique subpopulation identifier (specifying both locality and sampling timepoint) as a random effect to account for the temporal sampling structure within localities and the non-independence of individuals sampled at a given timepoint. To address the specific hypothesis that parental species would differ in Z:A ratios and nucleotide diversity, we ran the same models but with ancestry group as a predictor variable and subpopulation nested within population as a random effect. Model fits were assessed and confirmed using *DHARMa* (Hartig, 2026), and pairwise contrasts were assessed using *emmeans* (Lenth & Piaskowski, 2026). To assess temporal stability of Z:A ratios within populations, we conducted Wilcoxon rank-sum tests to compare each population at historical and contemporary sampling timepoints. To assess the effects of population contractions on Z:A, we compared population-level Z:A ratios in mainland vs. island populations using a Wilcoxon rank sum test. To investigate whether Z-linked diversity was disproportionately reduced when genome-wide nucleotide diversity was reduced, we fit a linear mixed effects model in *nlme* (Pinheiro et al., 2023) with the population-level Z:A ratio as the response variable, macrosomal nucleotide diversity as the predictor variable, and population ID as a random effect to account for repeated sampling of mainland transect populations. We fit the same model, but with Z-linked nucleotide diversity as a predictor variable, to assess whether associations with Z:A ratios were stronger with Z-linked loci than with macrosomal loci. All descriptive statistics are reported in the text as means ± 1 SD.

Finally, we assessed the degree to which geomorphometric variables related to archipelago evolution (O’Dea et al., 2026) predicted population Z:A ratios on the Bocas islands (see Supplementary Materials for details). We focused on variables that served as proxies for the magnitude of population contraction on each island (current area, area at time of isolation from mainland, and the integration of island size and time since isolation), the timing of the population contraction (isolation age), and degree of current spatial isolation (distance to mainland, proximity index, and proportion of land within a 1, 4, 16, and 64 km radius of a given island). Due to variable collinearity (Figure S3), we conducted principal components analysis on variables related to island size (Figure S4) and spatial isolation (Figure S5) and extracted the first principal components of each. We then conducted a multiple linear regression to assess correlations between Z:A ratios and island size, age, and spatial isolation.

## RESULTS

### Island populations derive from the mainland introgression zone

To graphically display genomic affinities among all populations, we provide an admixture plot at *K* = 2 clusters corresponding to genetic differentiation between the parental species *vitellinus* and *candei*, which dominates genomic structure across the entire system (Figure 2B). This plot, when considered with the plumage phenotypes of males, demonstrates how the mainland populations can be subdivided into four ancestry groups, comprising the two parental species, a core hybrid zone where extensive genomic admixture occurs and an introgression zone on the mainland, where all individuals are genomically like *candei* but have male plumage like *vitellinus*.

The origin of manakin populations on the Bocas del Toro Archipelago has been a matter of some interest. While island manakins closely resemble *vitellinus* in male plumage, several authors have provided morphological, behavioral, and limited genetic evidence that their affinities lie with introgression zone populations on the mainland (Wetmore, 1972; Billo, 2011; Barske et al. 2023; Chiver et al., 2026). However, Olson (1993) considered the island populations “relictual,” and Billo (2011) suggested that they could be the product of an ancient hybridization event (independent of the modern hybrid zone) or even be ancestral to the two parental species. In admixture analysis, all sampled island populations clustered with *candei* and the mainland introgression zone (Figure 2B), despite their close phenotypic resemblance to *vitellinus*. Thus, it seems indisputable that they are derived from the same hybridization processes that gave rise to mainland introgression zone populations.

### Z:A ratios vary across the mainland transect

We observed significant differences in overall nucleotide diversity between the two parental species, with *M. vitellinus* populations (populations 9.5–11) harboring higher macrosomal and Z-linked π than populations with predominantly *candei* ancestry (populations 2–7; Figure S6). Interestingly, macrosomal and Z-linked π was lowest in the introgression zone populations (Figure S6). One possible explanation is that introgression of *vitellinus* traits into *candei* was accompanied by selective sweeps that reduced genomic diversity.

The median population-level Z:A ratio across the mainland transect was 0.528 (mean = 0.528 ± 0.063; range = 0.421–0.662; Table S1). Correcting for male-biased mutation rate using a generalized avian mutation rate ratio (i.e., dividing by 1.11; Irwin, 2018), the median Z:A estimate decreased to 0.476. We detected significant variation in individual-level Z:A ratios across populations of the mainland transect according to ancestry group. Post-hoc Tukey’s tests revealed that parental *M. candei* populations had significantly higher Z:A ratios than all other ancestry groups: parental *M. vitellinus* (GLMM: z ratio = 3.52, p = 0.003), the hybrid zone (z ratio = 4.21, p = 0.0001), and the introgression zone (z ratio = 3.87, p = 0.0006). Pairwise comparisons between all mainland transect populations are represented in Figure 3 and Table S2. The median Z:A ratios of introgressed and hybrid populations were generally similar to parental populations of *vitellinus*, with the exception of population 4, which was more similar to parental *candei*, and population 7, which was especially low (Figure 3). Of note, population 4 experienced introgression of a male *vitellinus* plumage trait more recently than other introgression zone populations (Long et al. 2024; see Discussion). The location of population 7 on a narrow land bridge (Figure 2A) may cause it to behave more similarly to an island in terms of area constraints or gene flow restriction, leading to lower Z:A ratios (see below).

**Figure 3.**
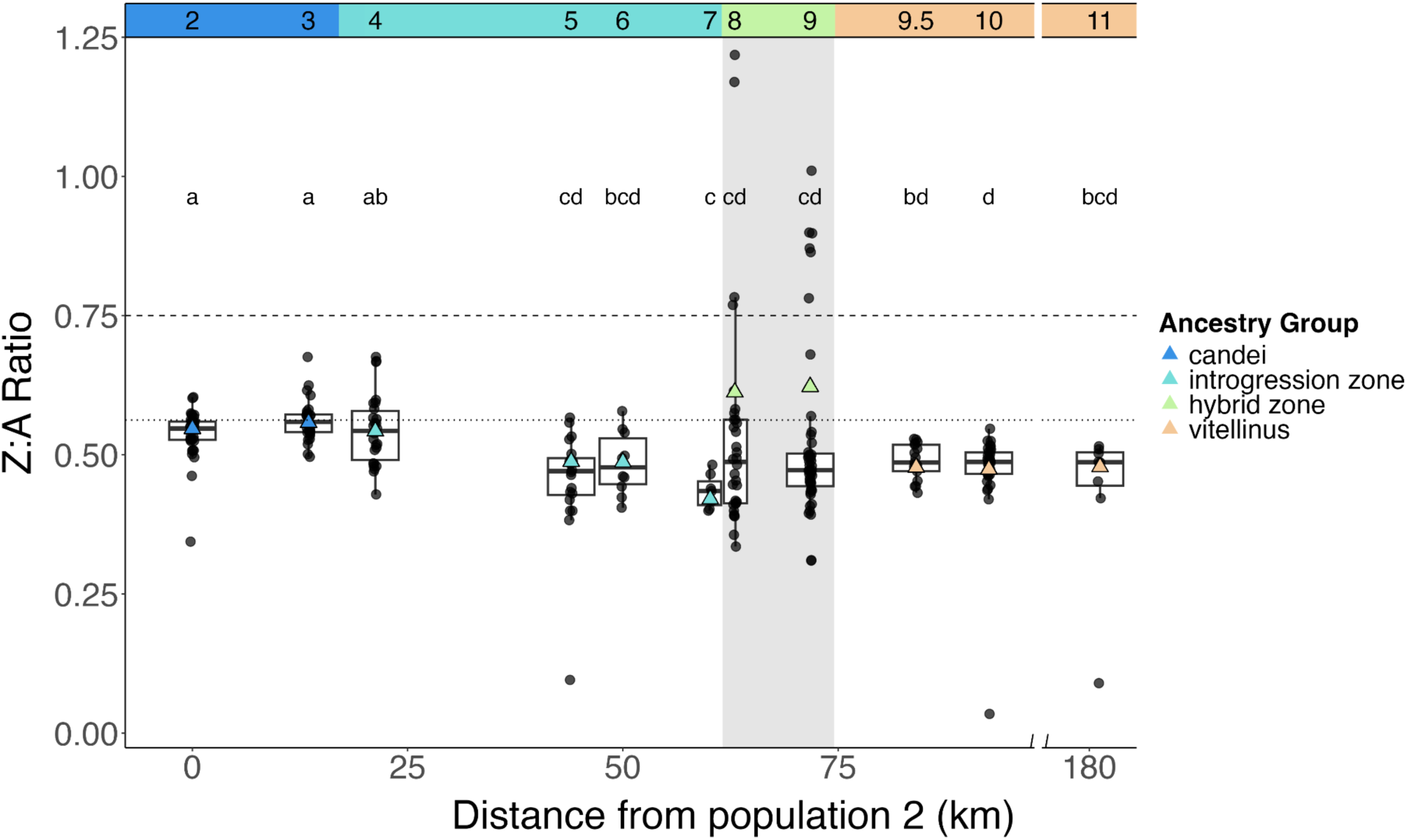
Across mainland populations, Z:A ratios are largely consistent within, but not between, ancestry groups. Populations are indicated by number and ancestry groups by color across the top of the graph. Triangles represent population-level point estimates of Z:A ratios calculated across all individuals in a population (calculations at each timepoint were averaged for populations that were sampled twice), while boxplots and circular points represent individual-level Z:A ratios. Lower-case letters above boxplots denote the results of Tukey’s post-hoc tests for pairwise significance; populations that share letters are not statistically different from one another based on the spread of individual-level Z:A values. Broadly, the three leftmost populations (the “a” group) differ statistically from most of the introgressed, hybrid, and *vitellinus* populations (the “c” and “d” groups). We did not observe any pairwise differences within ancestry groups along the transect with the exception of population 4, which differed from other introgressed populations and was similar to parental *candei* populations. We also note that the two populations that fall within the narrow genomic cline of the hybrid zone (populations 8 and 9, highlighted in gray) exhibited greater variance in individual-level Z:A estimates, as well as the highest average population-level Z:A estimates. The dashed horizontal line at y = 0.75 represents the theoretically expected population Z:A ratio under equilibrium conditions, while the dotted line at y = 0.5625 represents the minimum ratio attainable through male-biased reproductive skew alone.

In addition, variance in individual-level Z:A estimates increased at the hybrid zone center (populations 8 and 9), with some individuals exhibiting remarkably high Z:A ratios, a pattern not observed in any other mainland populations. Further inspection revealed that this pattern was driven by some hybrid individuals having unusually high Z-linked but not macrosomal diversity (Figures S6, S7). Given that the two parental species are more highly diverged on the Z chromosome than the macrosomes based on both *D_a_ and F_ST_* (Figure S8), this pattern likely arises from fast-Z effects in parentals leading to early-generation hybrids having disproportionately high Z-linked heterozygosity and thus higher Z:A ratios.

### Z:A ratios are temporally stable within mainland transect populations

Based on the distribution of individual Z:A ratios in mainland populations sampled at multiple timepoints, Z:A ratios were temporally stable within populations (Wilcoxon test: p > 0.05 for all within-population comparisons between timepoints; Figure 4). This was true even for populations 3 and 4, which experienced introgression of olive belly color between sampling timepoints. We note that the population-level point estimates of Z:A ratios in hybrid populations 8 and 9 fluctuated between historical and contemporary sampling timepoints (Figure 4), despite no statistically significant shifts occurring in the distribution of individual Z:A ratios. As the location of the genomic center of the hybrid zone was previously shown to be generally stable between sampling timepoints (Long et al. 2024), this pattern likely indicates the sensitivity of population-level calculations to a small number of individuals harboring high genetic diversity on the Z (e.g., recent hybrids; Figure S10).

**Figure 4.**
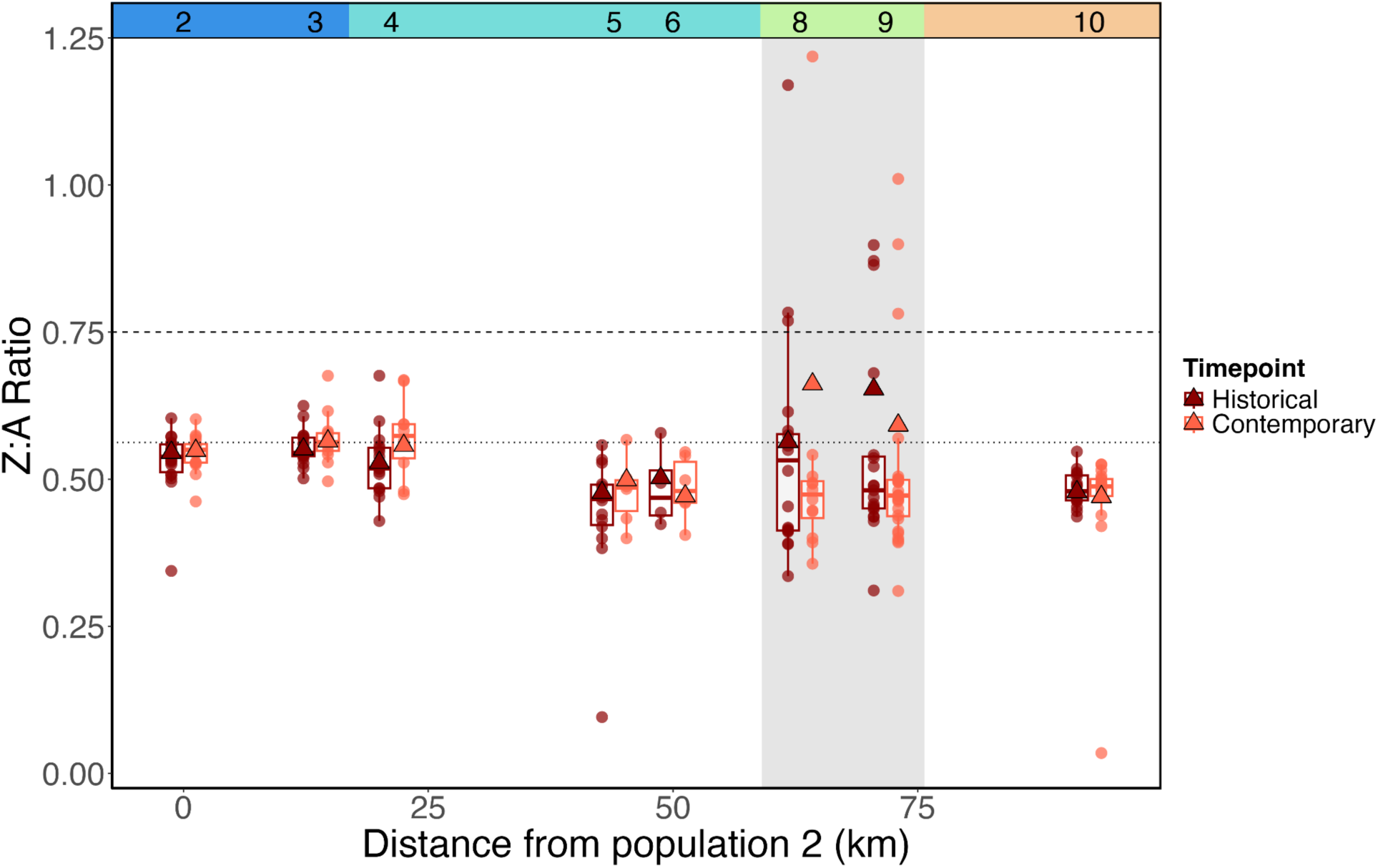
Z:A ratios are temporally stable within mainland populations. Z:A ratios were temporally stable within all mainland transect populations for which multiple sampling timepoints were available (p > 0.05 for all pairwise Wilcoxon tests, based on the values of individual-level Z:A ratios in populations at a given timepoint). Only mainland transect populations that were sampled at multiple timepoints are shown. Population IDs and ancestry groups are indicated across the top of the graph as in Figure 3. Triangles represent population-level Z:A ratios, calculated across all pairs of sequences in the population irrespective of individual, while boxplots and points represent the spread of Z:A ratios calculated within individuals. The dashed horizontal line at y = 0.75 represents the theoretically expected population Z:A ratio under equilibrium conditions, while the dotted horizontal line at 0.5625 represents the minimum Z:A ratio achievable through male-biased reproductive skew alone.

### Z:A ratios are reduced on islands

Macrosomal π was significantly lower in island populations than mainland populations (mean ± SD: 0.00085 ± 0.0002 vs. 0.00161 ± 0.0005, respectively; Wilcoxon rank sum: W = 0, p < 0.0001; Figure 5A). Similar patterns were observed for Z-linked π (mean ± SD: 0.0003 ± 0.0001 vs. 0.00086 ± 0.0003, respectively; Wilcoxon rank sum: W = 0, p < 0.0001; Figure 5B). Moreover, island populations exhibited significantly lower Z:A ratios than mainland populations (mean ± SD: 0.376 ± 0.077 vs. 0.528 ± 0.063, respectively; Wilcoxon: W = 9, p < 0.0001; Figure 5C). The median Z:A ratio among island populations after correcting for male mutation bias was estimated to be 0.337 (compared to 0.476 in mainland populations).

**Figure 5.**
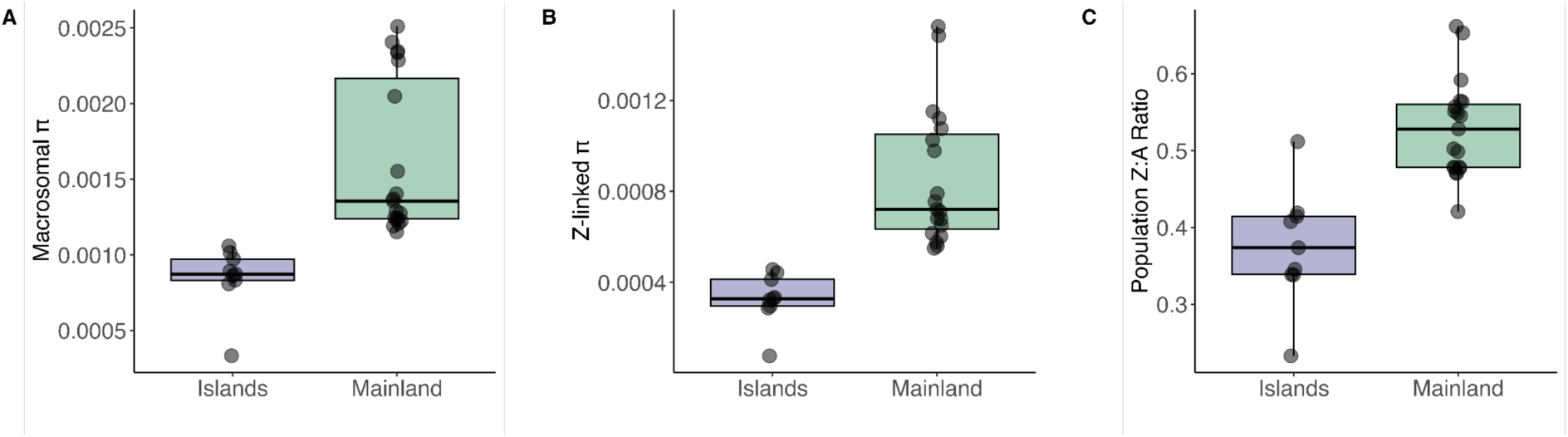
Island populations exhibit significantly lower macrosomal π, Z-linked π, and Z:A ratios than mainland populations. **(A)** Autosomal nucleotide diversity was significantly lower in island populations than mainland populations (W = 0, p < 0.0001). **(B)** Z-linked nucleotide diversity showed similar patterns (W = 0, p < 0.0001). **(C)** Z:A ratios were significantly lower in island populations than mainland populations (W = 9, p < 0.0001). This indicates that, while both macrosomal and Z-linked diversity were reduced on islands, losses in Z-linked diversity were disproportionate. Effects remained when islands were compared against only the introgression zone populations of *M. candei* (populations 4–7), which represent the closest mainland relatives of the island birds (macrosomal π: W = 0, p = 0.0002; Z-linked π: W = 0, p = 0.0002; Z:A ratio: W = 5, p = 0.003). Points represent values calculated at the population level.

Two island populations exhibited unusual Z:A ratios relative to other islands. The island of Escudo de Veraguas, which is the most spatially isolated and furthest from the mainland, had the lowest population Z:A ratio (∼0.23; Figure 5). At the other end of the spectrum, the small islet of Cayo Roldan exhibited unusually high Z:A ratios, more similar to those of mainland populations (Figure 5). Examining the spread of individual-level Z:A ratios within Roldan revealed a surprising degree of variation, a phenomenon not observed in any other islands (Figure S11). Given its proximity to the mainland, it is possible that individuals with high Z:A ratios on Roldan may be recent immigrants from the mainland, where Z-linked diversity is higher. We continue to explore geomorphometric predictors of Z:A ratios on Cayo Roldan and the other Bocas islands below.

### Declines in Z:A ratios are driven by declines in Z-linked diversity

Across all populations, Z:A ratios were positively associated with both macrosomal π (LMM: t = 3.23, df = 7, p = 0.01; Figure 6A) and Z-linked π (LMM: t = 6.13, df = 7, p < 0.001; Figure 6B). The fact that Z:A ratios decline even as macrosomal nucleotide diversity (the denominator) declines shows that reductions in overall genetic diversity must be associated with disproportionate reductions in Z-linked diversity. Indeed, correlations with the Z:A ratio across populations were stronger for Z-linked diversity (marginal R^2^ = 0.61; β = 0.86) than for macrosomal diversity (marginal R^2^ = 0.32; β = 0.57), further suggesting that reductions in the Z:A ratio are primarily due to dynamics on Z (Figures 6, S9). These effects were driven by the island populations, as well as the mainland populations with predominantly *M. candei* ancestry (populations 2–7; Figures 6C, 6D).

**Figure 6.**
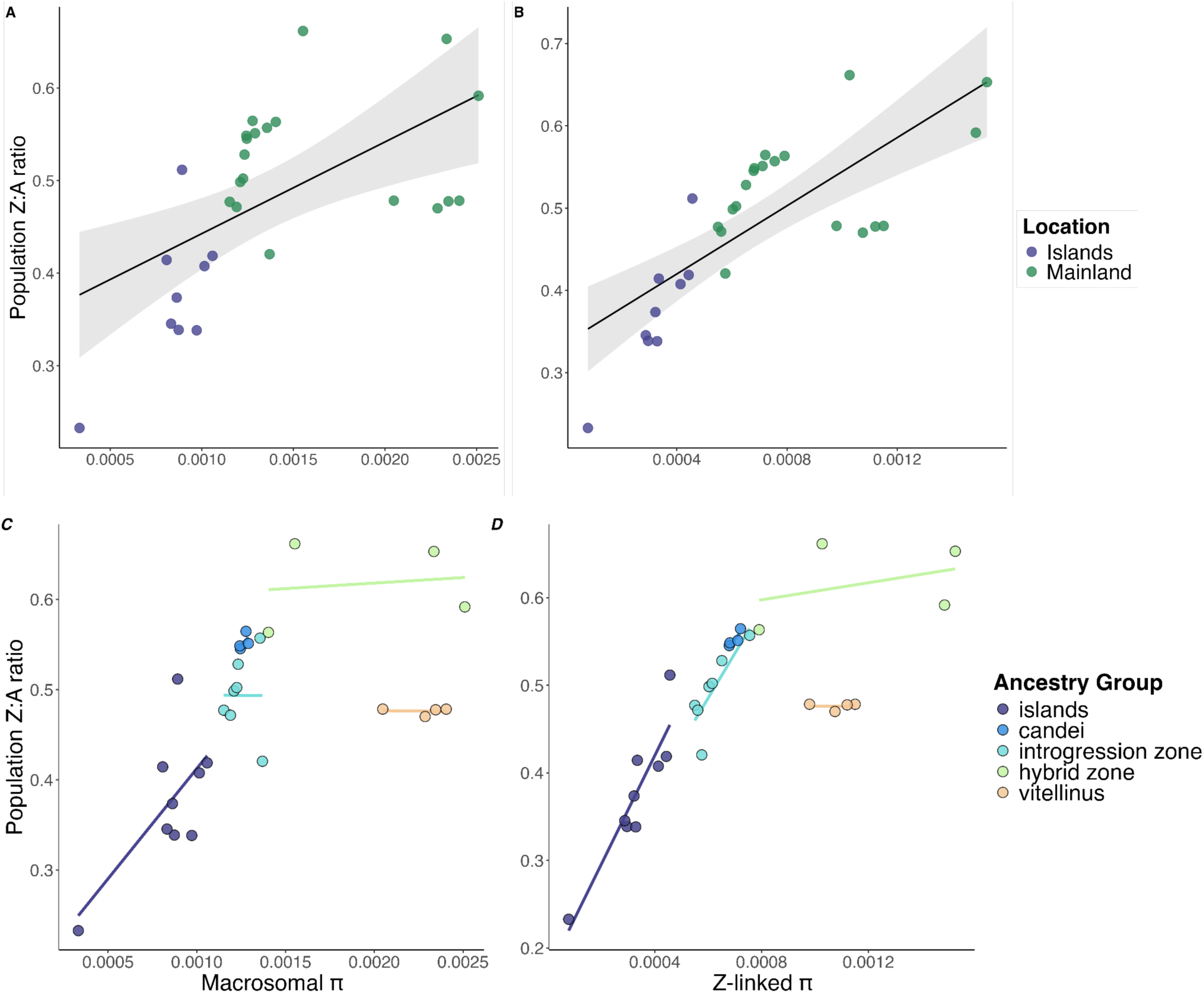
Reductions in the Z:A ratio are driven by disproportionate reductions of diversity on Z-linked loci relative to macrosomal loci. **(A)** Across all populations, Z:A ratios are positively associated with macrosomal diversity (LMM: t = 3.23, p = 0.01; marginal R^2^ = 0.32; β = 0.57), suggesting that Z-linked nucleotide diversity must be disproportionately affected when genome-wide nucleotide diversity is reduced. **(B)** Z:A ratios are more strongly associated with Z-linked diversity (t = 6.13, p < 0.001; marginal R^2^ = 0.61; β = 0.86) than with macrosomal diversity, again indicating that declines in Z:A are driven by dynamics on Z. Linear mixed effects models consider all populations together, with a random effect of population ID to account for repeated sampling of mainland populations. **(C, D)** Visualizing the trends within different ancestry groups reveals that correlations between the Z:A ratio and macrosomal diversity and Z-linked diversity are driven primarily by effects within the island populations and the mainland populations with predominantly *candei* ancestry.

### Spatial isolation is the strongest predictor of Z:A ratios on islands

We hypothesized that the magnitude of the population contraction (island size), the time since population contraction (island age), and gene flow (spatial proximity to other landmasses) would influence Z:A ratios on islands. Based on a multiple regression, neither island size PC1 (t = 0.13, p = 0.90) nor isolation age (t = –0.43, p = 0.68) significantly predicted Z:A ratios on islands (Table 1; Figure 7). However, spatial connectivity PC1 significantly predicted insular Z:A ratios, with more isolated islands exhibiting lower Z:A ratios (t = 3.50, p = 0.02; Table 1; Figure 7). This composite connectivity variable was more predictive of Z:A ratios than any of the individual spatial variables it comprises (Table S3; Figure S13). We include correlations between insular Z:A ratios and all relevant individual variables from O’Dea et al. (2026) in the Supplementary Materials (Table S3; Figure S13).

**Figure 7.**
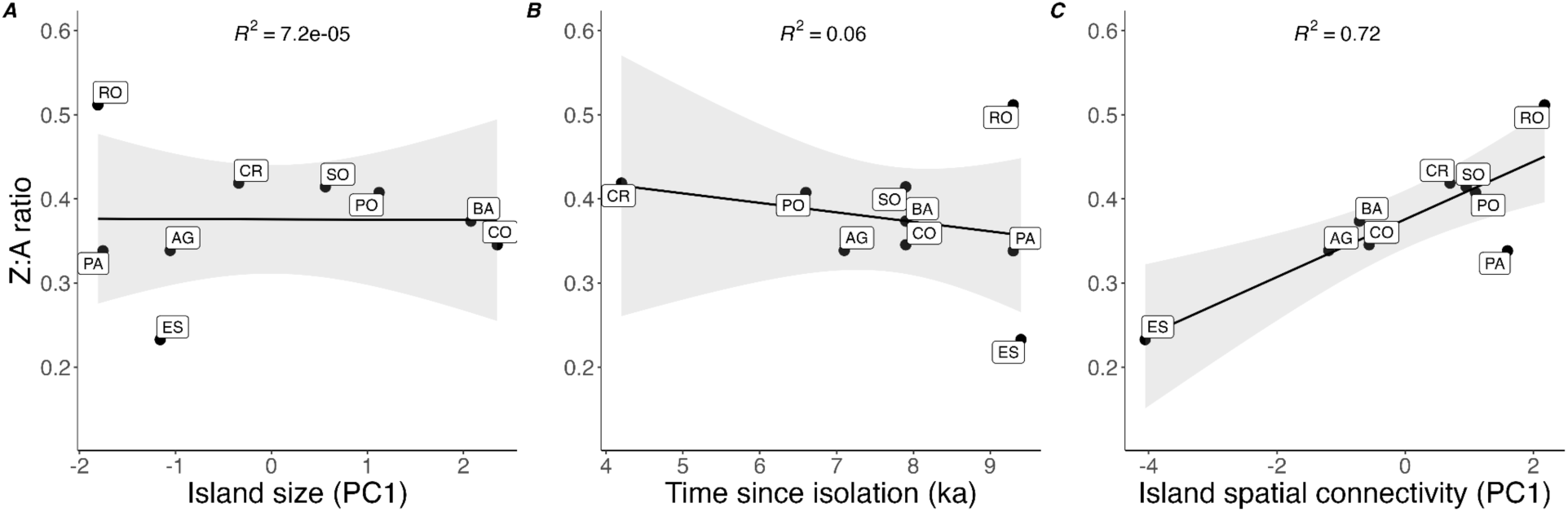
Island spatial connectivity, but not size or age, predicts population Z:A ratios in the Bocas del Toro Archipelago. **(A)** The first principal component (PC1) of variables related to island size (i.e., current area, area at time of isolation, and habitat-years) was not predictive of Z:A ratios on islands (LM: t = –0.02, p = 0.98; R^2^ < 0.0001). **(B)** Time since isolation (i.e., island age in ka) was also not predictive of Z:A ratios on islands (LM: t = –0.67, p = 0.53; R^2^ = 0.06). **(C)** The first principal component of variables related to island spatial connectivity (i.e., distance to mainland, proximity index, and B indices) significantly predicted Z:A ratios on islands (LM: t = 4.20, p = 0.004; R^2^ = 0.72).

**Table 1.** Output of multiple regression model assessing geomorphometric predictors of population Z:A ratios on islands of the Bocas del Toro Archipelago.

| Predictor | Estimate | SE | *t* value | *p* value |
| --- | --- | --- | --- | --- |
| Intercept | 0.41 | 0.09 | 4.60 | 0.006 ** |
| Size (PC1) | 0.001 | 0.01 | 0.13 | 0.90 |
| Age | −0.005 | 0.01 | −0.43 | 0.68 |
| Isolation (PC1) | **0.03** | **0.01** | **3.50** | **0.02 \*** |

When viewed as a function of overall spatial connectivity (i.e., potential for gene flow from neighboring landmasses, both mainland and island), the unusually high Z:A ratio for Cayo Roldan noted above is roughly in line with expectations for a highly spatially connected island (Figure 7). On the other hand, Isla Pastores, another small island near to the mainland, has low Z:A ratios for its degree of spatial connectivity (Figure 7), suggesting it may receive less immigration than expected. However, contrary to this hypothesis, Pastores has the lowest differentiation from the mainland introgression zone of any island population (Figure S12). Given their very small areas (22 and 201 ha, respectively), presumably small populations, and proximity to the mainland, we suggest that Z:A ratios on Roldan and Pastores may be especially variable due to the high potential for immigration and genetic drift to influence estimates.

## DISCUSSION

The Z:A ratio has been suggested as a potential genomic index of sexual selection strength in birds, with lower values indicating higher levels of male reproductive skew (Corl & Ellegren, 2012; Huang & Rabosky, 2015; Irwin, 2018; Schield et al., 2021). By leveraging longitudinal sampling data available in the bearded manakin system (genus *Manacus*), we empirically evaluated the degree to which this ratio fluctuates across known phylogenetic and biogeographic contexts. Our results indicate that Z:A ratios are temporally stable within mainland populations and vary between species in ways consistent with empirical data on variation in the intensity of sexual selection. However, processes other than sexual selection also have the potential to influence the Z:A ratio (Amster & Sella, 2020; Charlesworth, 2009; Ellegren, 2009a; Pool & Nielsen, 2007; Wilson Sayres, 2018), and the spatial variation in Z:A ratios we observed in the hybrid zone and island populations is consistent with expected effects of admixture and historical changes in population size on sex-linked genetic diversity.

Parental populations of *M. vitellinus* had lower Z:A ratios than parental populations of *M. candei*, which may indicate stronger sexual selection in the former. *M. vitellinus* exhibits several sexually selected plumage traits—notably, yellow collars and olive belly plumage—that are not present in *candei* but have introgressed asymmetrically, apparently under sexual selection, across a narrow hybrid zone (Parsons et al., 1993; Brumfield et al., 2001; Long et al., 2024). Prior research has shown that *vitellinus* and introgression zone males are more aggressive than *candei* males (McDonald et al., 2001), females prefer yellow-collared over white-collared males (Stein & Uy, 2006), and *vitellinus* produces more extreme versions of the physiologically challenging roll-snap display (Miles et al., 2018). Given these differences in traits relevant to sexual selection, as well as the expectation that potentially confounding life history variables (e.g., degree of sex-biased mutation, dispersal, generation times, etc.) should be similar between the two parental species, the lower Z:A ratios in *vitellinus* may indeed reflect stronger sexual selection operating in this lineage.

The observation that most introgression zone populations exhibit lower, *vitellinus*-like Z:A ratios (Figure 3) is consistent with the hypothesized scenario where plumage introgression is driven by sexual selection. The sole exception is population 4, where introgression occurred most recently, raising the possibility that Z:A ratios have yet to equilibrate in this population. Interestingly, and contrary to our expectations, the populations that underwent introgression of olive belly plumage between sampling timepoints (populations 3 and 4; see Figure 3E of Long et al. 2024) did not exhibit any detectable temporal shifts in Z:A ratios (Figure 4). Assuming that belly color and/or collar color have introgressed under sexual selection strong enough to reduce Z:A ratios, potential explanations for this result include: (1) introgression has occurred too recently to detect changes in Z:A ratios, or (2) comparisons between timepoints were underpowered to detect effects.

A notable perturbation in Z:A ratios occurred at the genomic center of the hybrid zone (Figures 3 and 4), where we observed increases in both individual-level variance and population-level Z:A ratios. This observation is likely attributable to “fast-Z” effects: due to higher evolutionary rates on the Z (Mank et al., 2007; Wanders et al., 2024), this chromosome exhibits higher between-lineage divergence than the rest of the genome (Figure S8). Recently admixed hybrids with one copy of each parental Z chromosome are then expected to have unusually high Z-linked *π* and Z:A ratios. In line with this interpretation, individuals with high Z:A ratios in the hybrid zone had unusually high Z-linked diversity (Figure S7) and were highly heterozygous for ancestrally informative parental alleles on the Z (Long et al., in prep). (Conversely, outlier individuals with very low Z:A ratios in other populations (Figures 3, 4 and S11) may be the result of close inbreeding resulting in high levels of homozygosity on Z.) Another potential contributor to higher population-level Z:A ratios in hybrid populations could be female-biased mortality, as expected under Haldane’s Rule, which posits that if one sex exhibits hybrid inviability or sterility, it is typically the heterogametic sex (in this case, the ZW females; Haldane, 1922). Female-biased mortality would produce higher Z:A ratios by disproportionately reducing autosomal diversity relative to Z-linked diversity. While there is evidence of reduced hatching success in this hybrid zone (Long et al., 2025), it is unknown whether inviability is sex-specific and thus likely to influence Z:A ratios. Because the overall distributions of Z:A ratios in hybrid populations closely resemble those in *vitellinus* and introgressed populations, apart from high Z-linked diversity in a few recently admixed individuals, we interpret fast-Z effects to be the more influential contributor to increased Z:A ratios in this hybrid zone.

Notably, we also observed a dramatic reduction in Z:A ratios in island populations relative to mainland populations (Figure 5), which was driven by disproportionate reductions in nucleotide diversity on the Z (Figure 6). While there are many examples of reduced genetic diversity on islands relative to the mainland (e.g., Frankham, 1997; Gyllenhaal et al., 2025; Whiteman et al., 2006), there have been fewer empirical demonstrations of island reductions of sex-linked genetic diversity relative to the autosomes. The islands of the Bocas del Toro Archipelago were formerly part of mainland Panama (O’Dea et al., 2026), and thus contemporary island populations presumably split from larger mainland populations primarily by vicariance. The inundation and formation of the archipelago would have reduced island population sizes (Runemark et al., 2012) resulting in losses of genetic diversity by drift. Coalescent modeling shows that population contractions should reduce diversity at Z-linked loci more rapidly than autosomal loci due to their smaller *N_e_*, producing transient reductions in Z:A ratios (Pool & Nielsen, 2007). Although Z-linked diversity should also rebound from bottlenecking events more rapidly than the autosomes (Pool & Nielsen, 2007), islands have little opportunity for subsequent population expansion after initial contraction due to area constraints; therefore, insular Z:A ratios are expected to remain relatively low on islands that were recently isolated, as is the case in the Bocas del Toro Archipelago. Moreover, weaker population size contractions (e.g., on the order of a reduction from 10,000 to 500 individuals) are expected to result in milder but longer-lasting reductions in the Z:A ratio (e.g., lasting > 5,000 generations), whereas severe population contractions (e.g., reductions from 10,000 to 20 individuals) are expected to result in stronger but shorter-lived reductions in Z:A (e.g., lasting ∼400 generations; Pool & Nielsen, 2007). Given the likelihood that island populations in this study were isolated between 2.9 and 9.5 ka (median = 8.1 ka; O’Dea et al., 2026), and yet Z:A ratios remain roughly 25% lower than mainland populations, a weaker population contraction is perhaps more probable. However, we note that dispersal scenarios and founder effects are also possible, particularly given the experimentally demonstrated and unusually strong ability (relative to other understory birds) of *Manacus* manakins to cross large water gaps (Moore et al., 2008). Such a scenario could help explain the high Z:A median and variance on Cayo Roldan (Figures 7, S11), a small island located just 1.3 km from the mainland.

Relatedly, we found that spatial isolation was the strongest predictor of Z:A ratios on the Bocas islands (Figure 7). Less spatially isolated islands likely receive higher rates of immigration from the mainland or other islands, both of which are expected to disproportionately contribute to Z-linked diversity over autosomal diversity: mainland immigrants are expected to have higher individual Z:A ratios, whereas immigrants from other islands are likely to be especially diverged on Z due to faster-Z effects. The positive correlation between spatial connectivity and Z:A ratios suggests that gene flow mitigates the effects of population contraction on insular Z-linked diversity, particularly given that proxies for the magnitude and timing of population contractions (i.e., island size and age) were not predictive. However, our area-related proxies may not represent the true magnitude of population contraction on islands, and therefore these conclusions should be considered tentative.

While the low Z-linked diversity on islands relative to the mainland may be predominantly attributable to their distinct demographic history, we note that differences in sexual selection strength on islands may also contribute. *Manacus* population density is noticeably higher on some islands compared to the mainland (M. J. Braun, personal observation), which has the potential to influence the degree of male-male competition and reproductive skew at leks (Hernandez, 1999). In addition, manakins on Escudo de Veraguas exhibit insular gigantism (Wetmore, 1959), a phenotype that could reflect the release of predator-related constraints on directional sexual selection for larger body size (Ponti et al., 2023). This scenario seems plausible, given that similar increases in male body size have been observed in lek-mating iguanas on the Galápagos (Wikelski & Trillmich, 1997), and body size has been found to be a positive predictor of male mating success in other species of *Manacus* (Shorey, 2002). A recent behavioral study found that the roll-snap display of Escudo males was similar to the slower rate observed in mainland *M. candei* (Chiver et al., 2026), which was interpreted to be inconsistent with the idea that sexual selection is stronger on this island; however, it is also possible that the evolution of gigantism in Escudo manakins constrained the maximum speed of the roll-snap, and thus this trait may not be directly comparable between mainland and island populations as an indicator of sexual selection strength. Regardless, it remains possible that demographic history and sexual selection differences jointly influence island Z:A ratios on islands, but continued research in insular populations is needed to better understand this interplay.

Finally, it is notable that most *Manacus* populations exhibited Z:A ratios that were not only lower than the Z:A ratio expected under neutrality (Z:A = 0.75), but also lower than the theoretical minimum achievable at equilibrium through male reproductive skew alone (Z:A = 0.5625; Charlesworth, 2001; Charlesworth 2009; Figure 1). While low Z:A ratios on islands are likely attributable to recent population contractions associated with island formation, factors beyond sexual selection must also be invoked to explain the very low mainland Z:A values. One possibility is that prior or ongoing demographic events (e.g., population contractions due to habitat loss) have led to losses in relative Z-linked diversity in mainland populations analogous to—but less dramatic than—the processes that have occurred on the islands. Indeed, anthropogenic deforestation and habitat loss is a rampant issue in western Panama. Selective sweeps on Z-linked genetic loci, which often harbor genes relevant to male sexual traits (Irwin, 2018), may also have acted to reduce the relative genetic diversity on the Z chromosome in this species complex. The *BCO2* locus involved in the production of the yellow collar phenotype that has introgressed from *vitellinus* to *candei* is located on a microsome (chromosome 24; Lim et al., 2024); given that our conclusions are based on Z:A ratios calculated relative to macrosomes, the presumed sweep associated with this introgression cannot itself be driving the Z:A patterns observed here, although the potential remains that unknown selective sweeps on Z could have influenced Z-linked diversity across populations. Moreover, we note that the drivers of the low Z:A ratios across populations must have counteracted the opposing effect of longer male:female generation time ratios in this system, which are expected to bias Z:A ratios *upward*: as the average reproductive age of males increases relative to females, Z-linked loci accumulate more mutations per generation due to ongoing male germline mutation throughout the lifespan, thereby increasing Z-linked polymorphism and Z:A ratios (Amster & Sella, 2016; 2020).

## CONCLUSIONS

The temporal stability of the Z:A ratio within mainland populations, as well as its variation in accordance with suspected differences in sexual selection between *Manacus* species, suggests its potential as a metric of sexual selection strength in scenarios where confounding non-equilibrium processes are largely controlled. However, given its apparent sensitivity to demographic processes such as admixture and changes in population size, we also advise caution when using the Z:A ratio as a metric of sexual selection in instances where the influence of these factors is unknown. We advocate for the use of approaches similar to those taken here—where comparisons are made among numerous closely related populations—as it allows for reasonable inference about the forces responsible for shaping Z-linked diversity that may not be possible in family-wide or single-population study designs. Incorporating coalescent-based methods for inferring ancient demographic events in tandem with Z:A ratio estimation (e.g., Chase et al., 2025) may also prove useful for disentangling the effects of confounding processes on sex-linked diversity when the primary aim is to isolate and infer the strength of sexual selection *per se*. As demonstrated here, the unique properties of the Z chromosome provide a window into complex demographic and selective processes, offering a promising avenue for illuminating the mechanisms that shape genetic diversity and evolutionary trajectories in natural populations.

## Data Availability

All supporting data and code to reproduce analyses will be made freely available on GitHub and/or Dryad following publication.

## Funding

National Science Foundation, Grant/Award Numbers: DEB 1457541 RCN, DGE 10-69157, and DGE 2139911; USDA National Institute of Food and Agriculture, Hatch project 1026333, Grant/Award Number: ILLU-875-984; Smithsonian Institution National Museum of Natural History; Tulane University; University of Maryland; University of Illinois Urbana-Champaign.

## Conflicts of Interest

The authors declare no conflicts of interest.

## Ethics Approval

All research was conducted in accordance with access, collecting and export permits provided by the Autoridad del Canal de Panamá, the Ministerio del Ambiente de Panama (MiAmbiente) and its predecessors (ANAM, INRENARE) and Smithsonian IACUC Protocol SI-22013.

## ACKNOWLEDGMENTS

We thank Storrs Olson, Charles Handley, Robb Brumfield, Travis Glenn, Judy Blake, Antonio Pineda, Sr, Antonio Pineda, Jr, Adolfo Muñoz, Ovidio Jaramillo, André Nguyen, Alexander Worm, Ivy Ciaburri, Olivia Ferrari, Elena Prado-Ragan, Vaughn Hage, Luis Ramos Vasquez, and Evalynn Trumbo for contributions to field work; Cesar Romero and the Chiriquí Land Company for hospitality in Bocas del Toro; the Jimenez, Lezcano, Aguilar, and Lopez families, as well as the Teribe and Ngäbe-Buglé people, for supporting our work; Owen McMillan, Lil Camacho, Isis Ochoa, Raineldo Urriola, and other staff of the Smithsonian Tropical Research Institute for logistical assistance; Christopher Balakrishnan, Jordan Karubian, Samridhi Chaturvedi, Alex Gunderson, and Renata Ribeiro for discussion of the project and/or comments on the manuscript; Stephen Parry for statistical consultation; Carrie Craig, Faridah Dahlan, Katie Murphy, and Mary Faith Flores for assistance in the laboratory; Matthew Kweskin for technical support with the computing cluster; and the NMNH collections staff, including Chris Milensky and Chris Huddleston. Genomic analyses were performed on the Smithsonian High-Performance Cluster (SI/HPC), Smithsonian Institution (https://doi.org/10.25572/SihPc).

## AUTHOR CONTRIBUTIONS

Conceptualization: HLA, KFPB, AO, TJP, MJB

Data curation: HLA, KFPB, KML

Formal analysis: HLA

Funding acquisition: HLA, JDB, MJB

Investigation: HLA, KFPB, KML, MJB

Methodology: HLA, KFPB, KML

Project Administration: HLA, MJB

Resources: JDB, AO, TJP, MJB

Supervision: MJB Validation: HLA, KFPB, KML

Visualization: HLA, KFPB Writing (original draft): HLA

Writing (review and editing): All authors

## Notes

### Competing Interest Statement

The authors have declared no competing interest.

