## Supplementary Materials for "Can the Z:A ratio serve as a genomic index of sexual selection? An assessment across space and time in a genus of lekking birds"

### Materials and Methods

#### *Sample collection and storage*

We obtained blood or tissue samples from *Manacus* manakins (n = 434) using ground-level mist nets or shotguns. Blood samples were obtained from live birds by collecting < 100 µL of blood per bird via brachial venipuncture; these birds were color-banded and released. Blood samples were stored in Longmire's lysis buffer (Longmire et al., 1997) in the field, then stored at –20°C before being transferred to –80°C freezers for long-term storage. Tissue samples (heart, liver and pectoral muscle) were obtained from birds that were prepared as museum specimens; these were frozen in liquid nitrogen in the field and later transferred to –80°C or –120°C for long term storage. All specimens and genetic samples are stored at the US National Museum of Natural History (Washington, DC). DNA was purified by phenol-chloroform extraction either manually or on a Gene Prep automated extractor (AutoGen, Inc.) at the Museum's molecular labs. The DNA samples have been used in a series of previously published or forthcoming studies (Bennett et al., 2025; Brumfield et al., 2001; Long et al., 2024; Parchman et al., 2013; Parsons et al., 1993; Vernasco et al., 2024; Yuri et al., 2009).

#### *RAD-seq library preparation*

Samples from most of the mainland transect populations appeared in Long et al. (2024), and thus these raw RAD-seq data were downloaded from the NCBI short-read archive. Libraries for other mainland and island population samples were prepared using similar laboratory protocols to Long et al. (2024), which was a modified version of the Etter et al. (2011) protocol. Briefly, DNA was digested using the restriction enzyme SbfI (New England BioLabs), barcoded and multiplexed, sheared and size-selected (fragment size 300-600 bp), adapter-ligated, and PCR-amplified. Sequencing was conducted by Azenta Life Sciences Group, Novogene Corp, the University of Illinois Urbana-Champaign Core Sequencing Facility, or the University of

Oregon Genomics & Cell Characterization Core Facility across 6 lanes on Illumina HiSeq 4000 or Illumina NovaSeq 6000 sequencing platforms in 150-bp paired-end reads.

*Bioinformatic processing*

We used process_radtags in *Stacks* v. 2.66 (Rochette et al., 2019) to demultiplex raw reads using custom inline barcodes and perform initial quality control. Specifically, we rescued barcodes and SbfI cut sites (--rescue) and discarded reads with uncalled bases (--clean), low quality scores (--quality), or average quality scores below Q15 (--score-limit 15) in a sliding window equal to 10% of the total read length (--window-size 0.1). Processed reads from all samples were then aligned to the *M. vitellinus* reference genome (NCBI RefSeq Accession: GCF_001715985.3) using *bwa* mem version 0.7.17 (Li, 2013; Li & Durbin, 2009).

We called SNPs and removed PCR duplicates using gstacks in *Stacks* v. 2.66. Based on the distribution of mean coverage across all samples, we implemented a minimum mean coverage threshold of 6x, resulting in 43 individuals (9.9%) being dropped. We also removed an individual with site missingness frequency 1–2 orders of magnitude higher than the other samples. With this final subset (n = 390 individuals), we conducted iterative filtering in *Stacks* populations to determine the individual and population missingness filters that optimized the tradeoff between stringency and site retention. In the final filtering scheme, sites needed to be present in all populations (-p 28, considering the same localities sampled at multiple timepoints as distinct populations) and at least 80 percent of the individuals in each population (-r 0.8). We also implemented a minimum minor allele count filter of 3 (-mac 3). Finally, we generated all-sites VCFs using *Stacks* populations (--vcf-all) for downstream processing in *pixy* (Korunes & Samuk, 2021).

On average, we retained approximately 14 individuals per population per timepoint (mean ± SD: 13.93 ± 7.02 individuals; range: 4–33) and 19.5 million total sites (mean ± SD: 19,545,876.61 ± 8,155.28 sites; range = 19,525,188–19,554,419). These values corresponded to roughly 91 percent of the total unfiltered individuals per population (mean ± SD: 90.85 ± 12.66%; range: 54.45–100%) and 24 percent of the total unfiltered sites per population (mean ± SD: 23.62 ± 5.96%; range: 12.06–31.62%). Of the total retained sites, roughly 18 million mapped to autosomes or the Z (mean ± SD: 18,803,490 ± 7,911.34 sites; range: 18,783,275–18,811,642); unplaced sites were excluded from analyses.

*Calculating individual-level Z:A ratios*

To obtain estimates of measurement uncertainty for statistical tests of spatial and temporal variation in macrosomal $\pi$, Z-linked $\pi$, and Z:A ratios across the mainland transect, we calculated diversity measures at the individual level, similar to individual heterozygosity. We opted to use individual-level measurements for this purpose—rather than chromosome-level measurements, as in Balakrishnan et al. (2026)—because (1) the autosomes are best treated as a single compartment in this case, given that the Z is expected to exhibit ¾ the nucleotide diversity of the autosomes on average, but selective events on individual chromosomes could skew chromosome-level ratios; and (2) individuals can be thought of as independent draws of entire genomic compartments (i.e., the Z chromosome vs the autosomes) from the population gene pool. In addition, individual-level measurements had the additional benefit of revealing the degree to which individual-level variability (particularly in the hybrid zone) influenced population-level Z:A estimates, which would not have been revealed with a bootstrapping approach.

*Assessing biogeographic predictors of Z:A ratios on islands*

We aimed to assess the relative influence of various biogeographic variables on island Z:A ratios, hypothesizing that the timing and magnitude of population contraction, as well as rates of immigration from the mainland and other islands, would be the primary factors influencing Z:A ratios on islands. We obtained geomorphometric data for the islands of the Bocas del Toro Archipelago from O'Dea et al. (2026), which used high-resolution bathymetry and historical sea-level data to reconstruct the geological history of the archipelago and estimate a suite of contemporary and historical metrics that could be informative about current and past population dynamics. As a proxy for the magnitude of the population contraction on each island, we considered variables from O'Dea et al. (2026) expected to correlate with population size since island formation: current area (ha), isolation area (i.e., the size of the island at the time of separation from the mainland), and habitat-years (i.e., the total area available annually since isolation, defined as the integration of island size vs. time since isolation, in ha/ka). Island isolation age (ka) was used as a measure of the time elapsed since population contraction. We also considered a suite of variables related to spatial isolation: distance to the mainland, proximity index, and the B1–B64 indices (defined below). Distance to the mainland was defined as the shortest straight-line distance (m) from a given island to the nearest mainland location. The proximity index was defined as the log-transformed sum of the areas of other islands, each inversely weighted by their distance to the focal island (i.e., $\log_{10}[\Sigma(A_i/d_i^2)]$, where $A$ = area and $d$ = distance of a given island; per Kalmar & Currie, 2006); this can be considered a measure of "stepping stone" connectivity to other islands. The B1, B4, B16, and B64 indices correspond to the proportion of land within 1, 4, 16, and 64 km of a given island, and indicate local, intermediate, sub-regional, and regional connectivity, respectively. The remaining variables from O'Dea et al. (decay rate, maximum elevation, and composite variables) were not considered because they were not directly related to any *a priori* hypotheses about Z:A ratios on islands. Due to the large number of conceptually related and collinear predictor variables (Figure S3), we conducted principal components analysis (PCA) on variables related to island size (Figure S4) and spatial isolation (Figure S5). PCAs yielded readily interpretable first principal components, and these were extracted and used for hypothesis tests related to population size and connectivity. We conducted a multiple linear regression using population-level Z:A ratios (calculated using macrosomal $\pi$ in the denominator) as the response variable, and island size (i.e., PC1 of historical and current size-related variables), isolation age, and spatial connectivity (i.e., PC1 of spatial variables) as predictors. Results were similar when Z:A ratios were regressed on each of the three predictor variables individually, and thus we present only the multiple regression results.

### Tables and Figures

**Table S1. Summary of post-filtering RAD-seq data used in analyses.** Z:A ratios are calculated using macrosomal nucleotide diversity (π) in the denominator (* = values corrected for male-biased mutation). Major Ancestry denotes the predominant ancestry component based on the ADMIXTURE analysis in Figure 2B.

| Population | Timepoint | Location | Major Ancestry | n | # Z-linked sites | # Macrosomal sites | $\pi_Z$ | $\pi_{macrosomes}$ | Z:A ratio | Z:A ratio* |
|---|---|---|---|---|---|---|---|---|---|---|
| 2 | Historical | Mainland | M. candei | 18 | 1,006,471 | 12,572,440 | 0.00068 | 0.00124 | 0.545 | 0.491 |
| 2 | Modern | Mainland | M. candei | 18 | 1,006,857 | 12,577,752 | 0.00068 | 0.00124 | 0.548 | 0.494 |
| 3 | Historical | Mainland | M. candei | 18 | 1,006,329 | 12,571,281 | 0.00071 | 0.00129 | 0.551 | 0.497 |
| 3 | Modern | Mainland | M. candei | 13 | 1,006,844 | 12,578,027 | 0.00072 | 0.00128 | 0.565 | 0.509 |
| 4 | Historical | Mainland | M. candei (introgressed) | 16 | 1,006,288 | 12,569,405 | 0.00065 | 0.00123 | 0.528 | 0.476 |
| 4 | Modern | Mainland | M. candei (introgressed) | 10 | 1,006,804 | 12,577,486 | 0.00076 | 0.00136 | 0.557 | 0.502 |
| 5 | Historical | Mainland | M. candei (introgressed) | 14 | 1,006,876 | 12,578,692 | 0.00055 | 0.00115 | 0.477 | 0.430 |
| 5 | Modern | Mainland | M. candei (introgressed) | 6 | 1,006,660 | 12,575,046 | 0.00060 | 0.00121 | 0.499 | 0.449 |
| 6 | Historical | Mainland | M. candei (introgressed) | 4 | 1,006,541 | 12,573,549 | 0.00062 | 0.00122 | 0.502 | 0.452 |
| 6 | Modern | Mainland | M. candei (introgressed) | 6 | 1,006,679 | 12,575,606 | 0.00056 | 0.00119 | 0.472 | 0.425 |
| 7 | Historical | Mainland | M. candei (introgressed) | 7 | 1,005,768 | 12,562,889 | 0.00058 | 0.00137 | 0.421 | 0.379 |
| 8 | Historical | Mainland | Hybrid | 20 | 1,006,998 | 12,580,436 | 0.00079 | 0.00140 | 0.563 | 0.508 |
| 8 | Modern | Mainland | Hybrid | 12 | 1,006,883 | 12,578,549 | 0.00103 | 0.00155 | 0.662 | 0.596 |
| 9 | Historical | Mainland | Hybrid | 19 | 1,006,989 | 12,580,551 | 0.00153 | 0.00234 | 0.653 | 0.588 |
| 9 | Modern | Mainland | Hybrid | 28 | 1,007,038 | 12,581,176 | 0.00149 | 0.00251 | 0.592 | 0.533 |
| 9.5 | Modern | Mainland | M. vitellinus | 17 | 1,006,654 | 12,575,439 | 0.00112 | 0.00235 | 0.478 | 0.430 |
| 10 | Historical | Mainland | M. vitellinus | 16 | 1,006,949 | 12,579,328 | 0.00098 | 0.00205 | 0.478 | 0.431 |
| 10 | Modern | Mainland | M. vitellinus | 18 | 1,006,853 | 12,577,868 | 0.00107 | 0.00229 | 0.470 | 0.424 |
| 11 | Historical | Mainland | M. vitellinus | 8 | 1,005,810 | 12,562,906 | 0.00115 | 0.00241 | 0.478 | 0.431 |
| Agua | -- | Islands | M. candei (introgressed) | 12 | 1,007,107 | 12,582,956 | 0.00030 | 0.00087 | 0.339 | 0.305 |
| Bastimentos | -- | Islands | M. candei (introgressed) | 33 | 1,007,120 | 12,583,249 | 0.00032 | 0.00086 | 0.374 | 0.337 |
| Cristóbal | -- | Islands | M. candei (introgressed) | 11 | 1,007,092 | 12,582,912 | 0.00044 | 0.00106 | 0.419 | 0.377 |
| Colón | -- | Islands | M. candei (introgressed) | 23 | 1,007,120 | 12,583,166 | 0.00029 | 0.00083 | 0.345 | 0.311 |
| Escudo | -- | Islands | M. candei (introgressed) | 14 | 1,006,959 | 12,581,582 | 0.00008 | 0.00033 | 0.233 | 0.210 |
| Pastores | -- | Islands | M. candei (introgressed) | 4 | 1,006,971 | 12,580,604 | 0.00033 | 0.00097 | 0.338 | 0.305 |
| Roldan | -- | Islands | M. candei (introgressed) | 6 | 1,007,095 | 12,582,488 | 0.00046 | 0.00089 | 0.512 | 0.461 |
| Solarte | -- | Islands | M. candei (introgressed) | 9 | 1,007,106 | 12,582,935 | 0.00033 | 0.00081 | 0.414 | 0.373 |
| Popa | -- | Islands | M. candei (introgressed) | 10 | 1,007,076 | 12,582,689 | 0.00041 | 0.00101 | 0.408 | 0.367 |

**Table S2. Significant pairwise post-hoc comparisons of populations along the mainland transect.** Values in the table represent the output of Tukey's post-hoc tests based on a generalized linear mixed effects model fit in *glmmTMB*. This model included individual-level Z:A ratios as the response variable, population (sampling locality) as a fixed effect, sampling timepoint within sampling locality as a random intercept, and a *t* error distribution to account for heavy tails. The Tukey-adjusted p values here and used to denote pairwise significance in Figure 3.

| Population A | Population B | Estimate | SE | z ratio | Adjusted p |
|---|---|---|---|---|---|
| 2 | 5 | 0.07 | 0.01 | 5.68 | 0.000 |
| 2 | 6 | 0.07 | 0.02 | 3.60 | 0.014 |
| 2 | 7 | 0.11 | 0.02 | 7.18 | 0.000 |
| 2 | 8 | 0.09 | 0.02 | 4.68 | 0.000 |
| 2 | 9 | 0.07 | 0.01 | 8.37 | 0.000 |
| 2 | 9.5 | 0.05 | 0.01 | 4.67 | 0.000 |
| 3 | 5 | 0.08 | 0.01 | 6.42 | 0.000 |
| 3 | 6 | 0.08 | 0.02 | 4.14 | 0.002 |
| 3 | 7 | 0.12 | 0.02 | 7.81 | 0.000 |
| 3 | 8 | 0.10 | 0.02 | 5.25 | 0.000 |
| 3 | 9 | 0.09 | 0.01 | 9.32 | 0.000 |
| 3 | 9.5 | 0.06 | 0.01 | 5.52 | 0.000 |
| 4 | 5 | 0.07 | 0.02 | 4.13 | 0.002 |
| 4 | 7 | 0.11 | 0.02 | 5.77 | 0.000 |
| 4 | 8 | 0.08 | 0.02 | 3.83 | 0.006 |
| 4 | 9 | 0.07 | 0.01 | 5.20 | 0.000 |
| 7 | 9.5 | -0.06 | 0.02 | -3.42 | 0.026 |
| 10 | 2 | -0.06 | 0.01 | -6.89 | 0.000 |
| 10 | 3 | -0.07 | 0.01 | -7.91 | 0.000 |
| 10 | 4 | -0.05 | 0.01 | -4.07 | 0.002 |
| 10 | 7 | 0.05 | 0.02 | 3.42 | 0.026 |
| 11 | 2 | -0.06 | 0.01 | -3.71 | 0.009 |
| 11 | 3 | -0.07 | 0.02 | -4.39 | 0.001 |

**Table S3.** Outputs of univariate regression analyses for Z:A ratios vs. relevant predictor variables from O'Dea et al. (2026). Data and regressions are visualized in Figure S13.

| Predictors | Proxy for: | Estimate | SE | t value | R-squared value | p value | Adjusted p value |
|---|---|---|---|---|---|---|---|
| Area (ha) | Current population size | 0 | 0 | 0.07 | 0.001 | 0.94 | 1 |
| Isolation area (ha) | Population size at time of isolation | 0 | 0 | -0.18 | 0.004 | 0.86 | 1 |
| Habitat-years (ha/ka) | Population size through time | 0 | 0 | 0.04 | 0 | 0.97 | 1 |
| Time since isolation (ka) | Time since contraction | -0.011 | 0.017 | -0.67 | 0.06 | 0.53 | 1 |
| Distance to mainland (km) | Potential for mainland gene flow | -0.009 | 0.004 | -2.29 | 0.428 | 0.06 | 0.39 |
| Proximity index | Potential for inter-island gene flow | 0.035 | 0.013 | 2.61 | 0.492 | 0.04 | 0.3 |
| B1 index | Potential for gene flow | 1.726 | 1.048 | 1.65 | 0.279 | 0.14 | 0.72 |
| B4 index | Potential for gene flow | 0.514 | 0.193 | 2.67 | 0.504 | 0.03 | 0.3 |
| B16 index | Potential for gene flow | 0.217 | 0.109 | 1.99 | 0.362 | 0.09 | 0.52 |
| B64 index | Potential for gene flow | 1.027 | 0.379 | 2.71 | 0.512 | 0.03 | 0.3 |

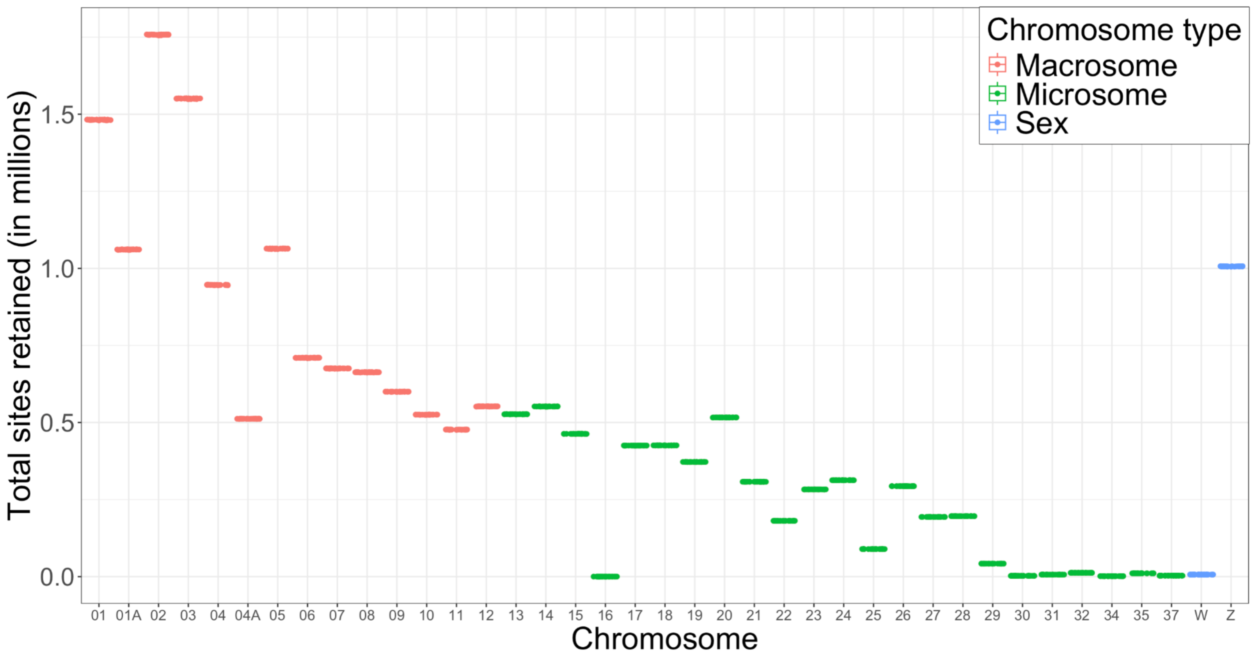


**Figure S1. Number of sequenced sites per chromosome retained after filtering.** Each point represents the total number sites per population at a given sampling locality. Due to relatively strict filtering, all populations retain similar numbers of sites on each chromosome; see Methods for operational definition of chromosomes. Sites that could not be associated with a chromosome were not used in any analyses and are not depicted here.

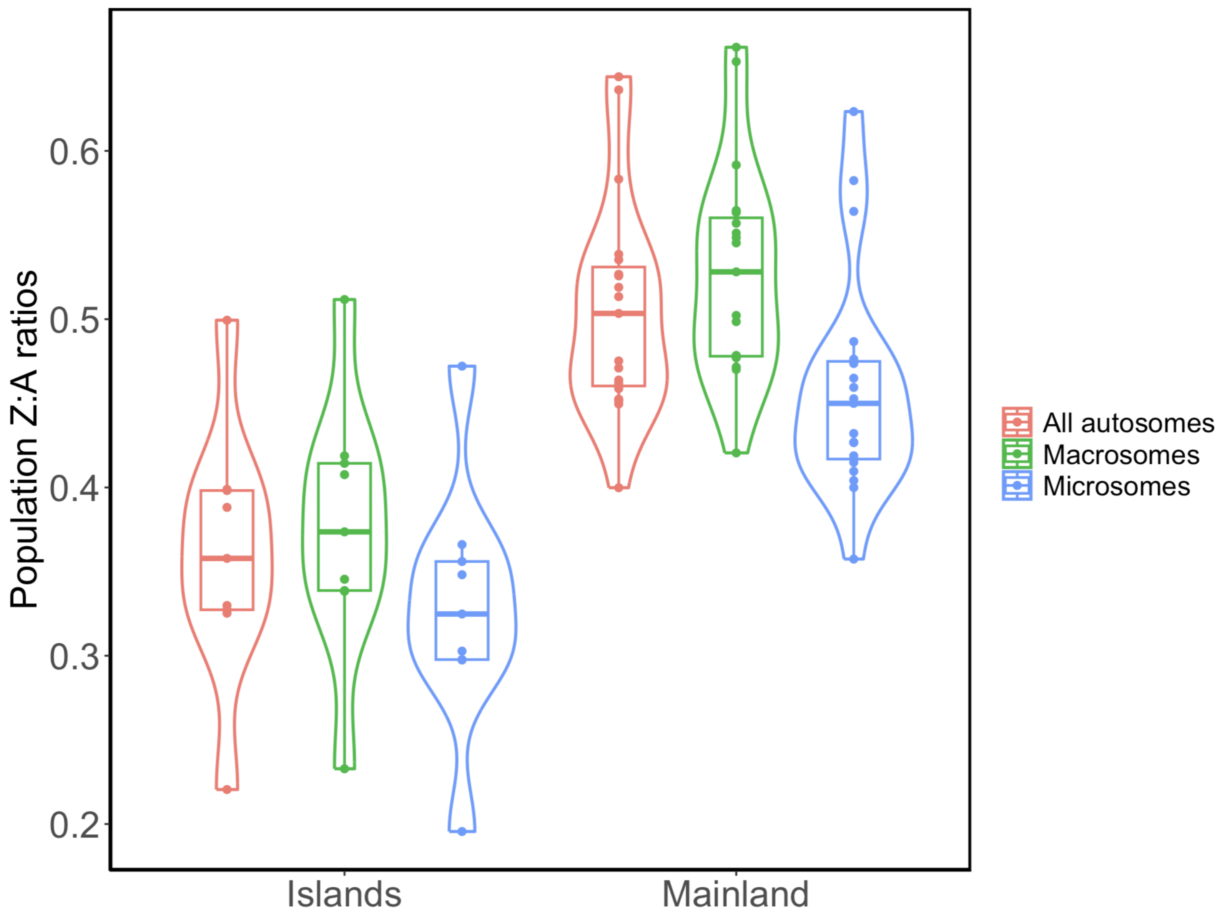


**Figure S2. Differences in population-level Z:A estimates when all autosomes, only macrosomes, or only microsomes are used as the denominator in Z:A calculations.** Z:A estimates were significantly lower when using microsomal vs. macrosomal $\pi$ as the denominator value, likely due to the higher recombination rates of microsomes compared to larger chromosomes resulting in higher nucleotide diversity on microsomes and lower Z:A ratios. See Methods for operational definitions of chromosomes, macrosomes and microsomes.

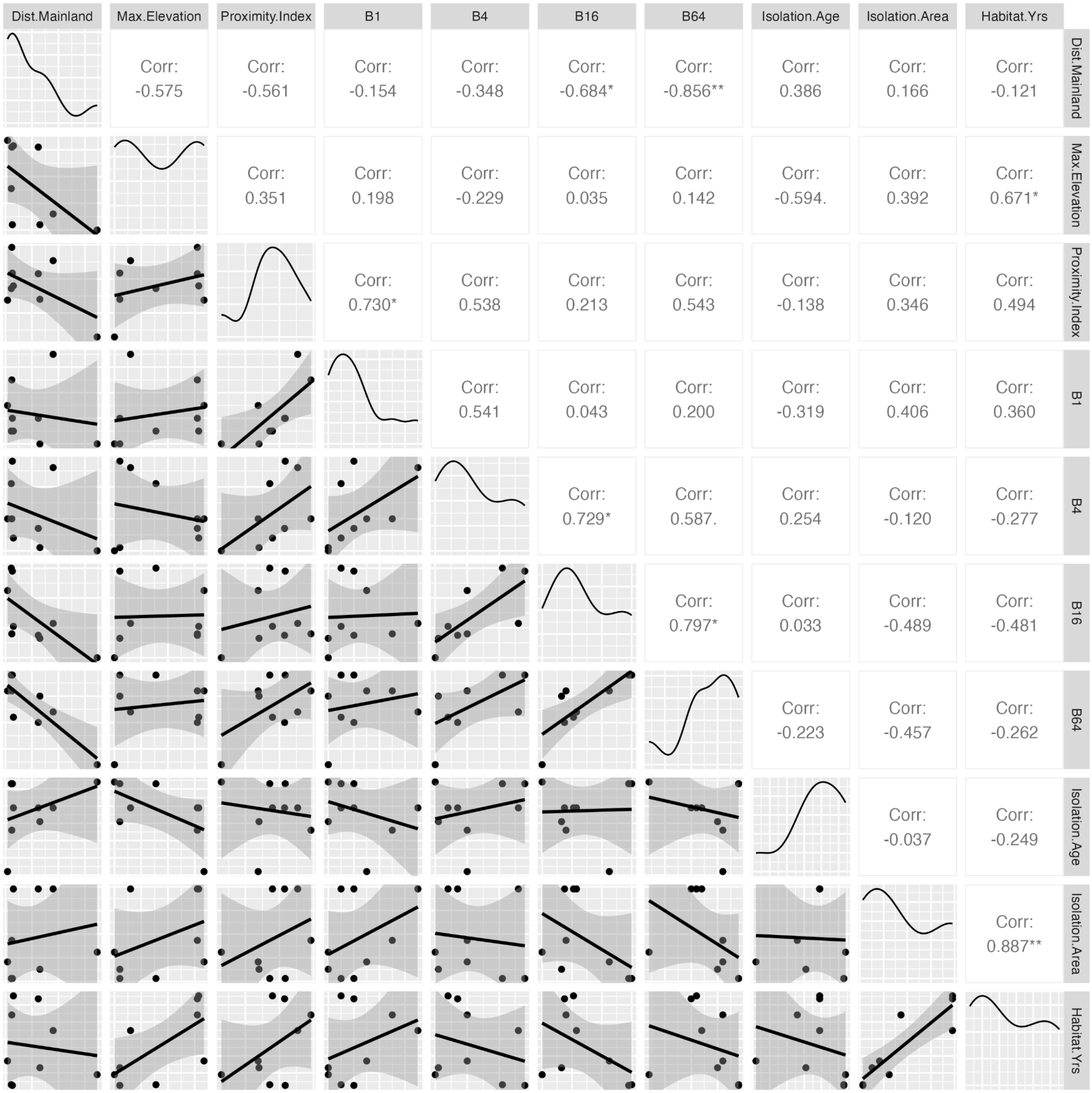


**Figure S3. Pairwise correlations between geomorphometric variables from O'Dea et al. (2026) potentially relevant to island Z:A ratios.** Predictor variables fall into three major classes: variables related to island size (current area, isolation area, and habitat-years); variables related to spatial isolation (distance to mainland, proximity index, and B indices); and variables related to time since island formation (isolation age). Pearson's correlation coefficients of pairwise comparisons are shown, and significant correlations are denoted by asterisks (* $p < 0.05$; ** $p < 0.01$).

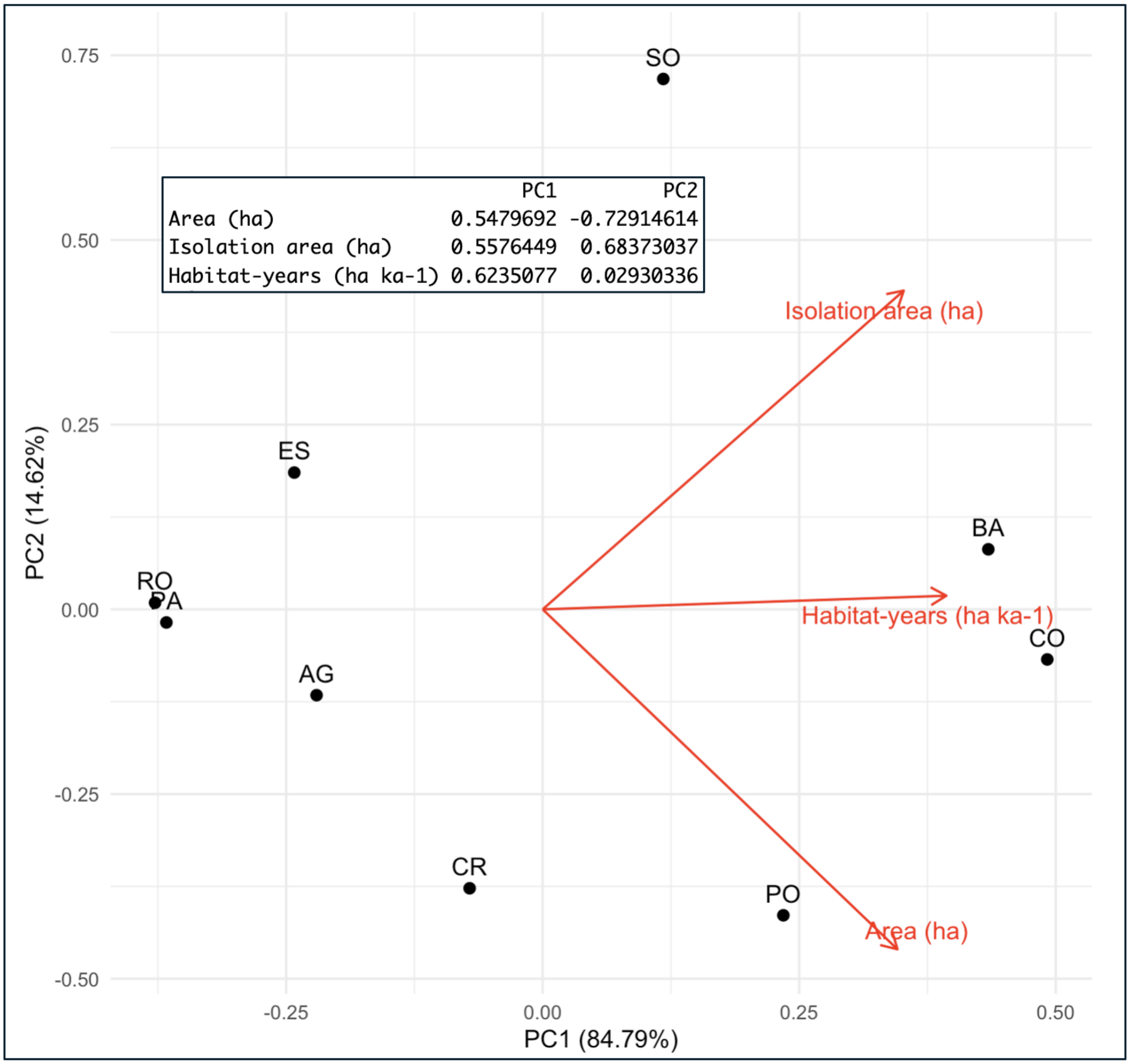


**Figure S4. PCA of variables related to modern and historical island size.** All area-related variables load positively along PC1. As such, islands with high values of PC1 have had greater available area (and likely greater *Manacus* population sizes) since their isolation, while islands with low values of PC1 have been smaller throughout their history. Inset: loadings for PC1 and PC2.

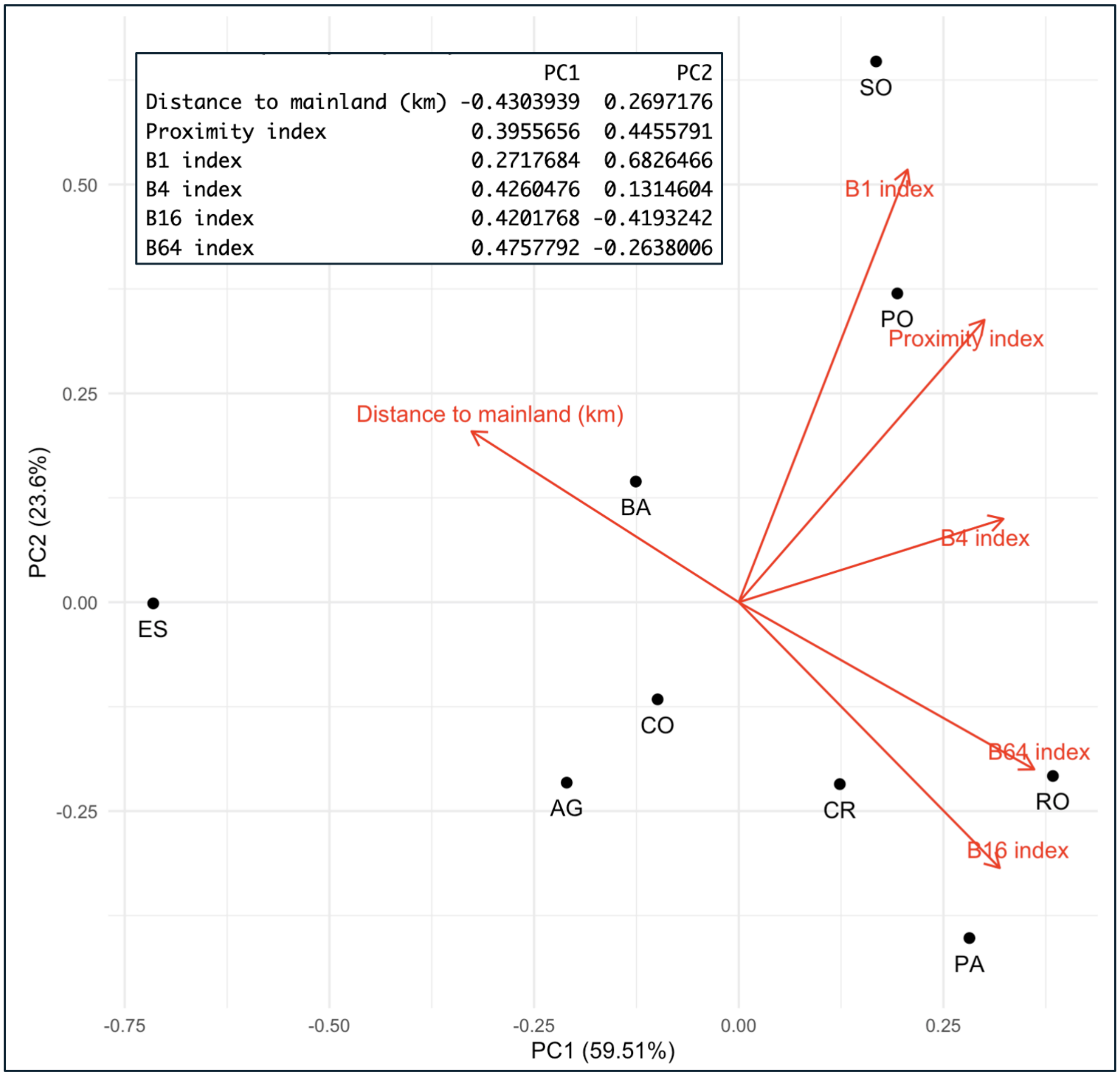


**Figure S5. PCA of variables related to island spatial isolation.** Variables related to proximity of other landmasses (proximity index, B indices) load positively along PC1, while distance to the mainland loads negatively. As such, variables with high values on PC1 are more spatially connected to other landmasses, while variables with low values on PC1 are more spatially isolated from other landmasses. Inset: loadings for PC1 and PC2.

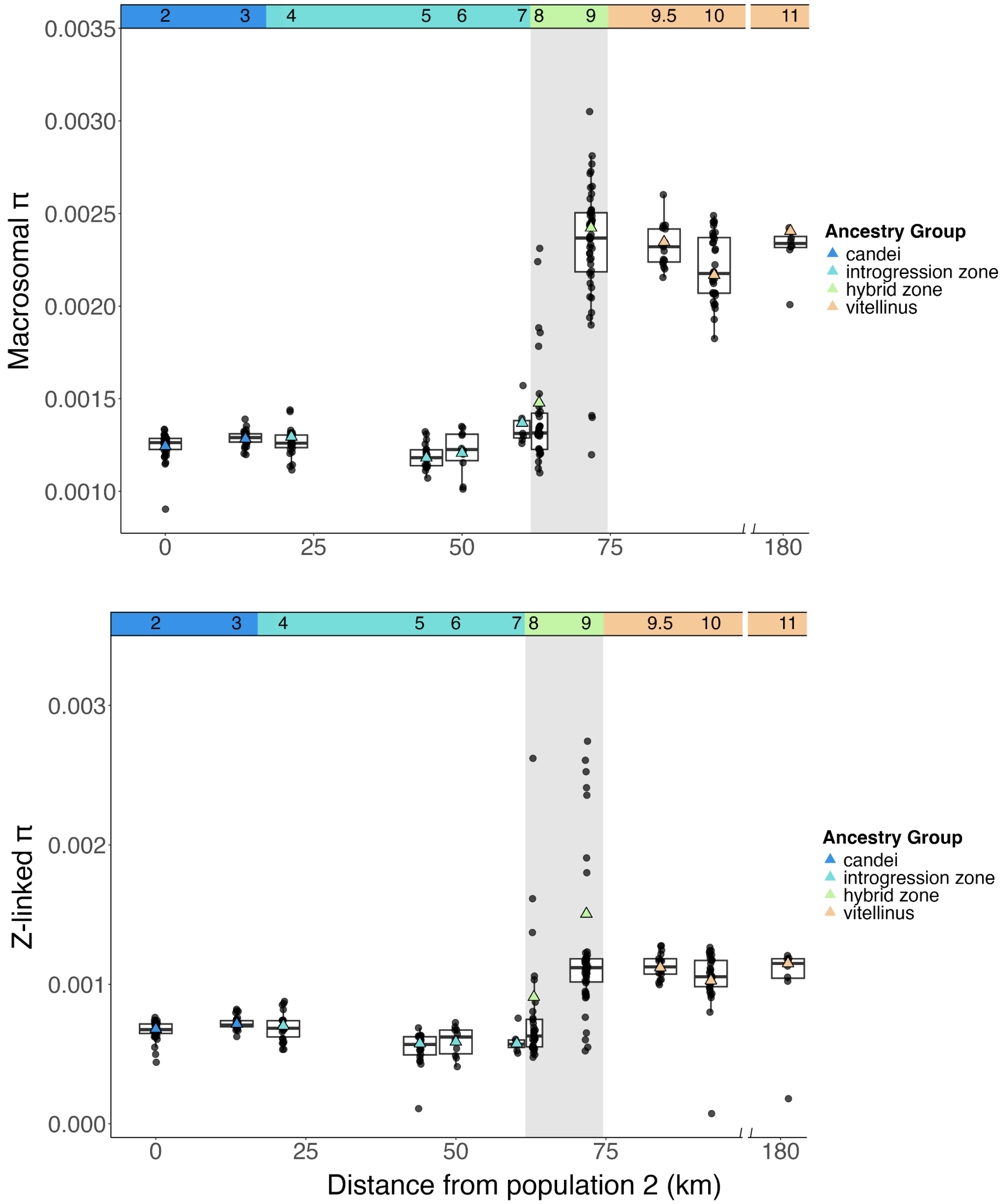


**Figure S6. Macrosomal and Z-linked nucleotide diversity across the mainland transect. (A)** *M. vitellinus* populations had significantly higher macrosomal π than introgression zone (GLMM: z ratio = 5.71, $p < 0.0001$) and parental populations of *M. candei* (z ratio = 4.78, $p < 0.0001$). **(B)** *M. vitellinus* populations had higher Z-linked π than introgression zone (GLMM: z ratio = 5.50, $p < 0.0001$) and parental populations of *M. candei* (z ratio = 3.94, $p = 0.0005$). Points represent individual-level nucleotide diversity for (A) macrosomes and (B) Z chromosomes in each of 11 mainland transect populations. Individuals sampled in both historical and contemporary timepoints are represented. Triangles represent population-level nucleotide diversity; for populations that were sampled twice, the population-level nucleotide diversity values calculated for historical and contemporary timepoints were averaged.

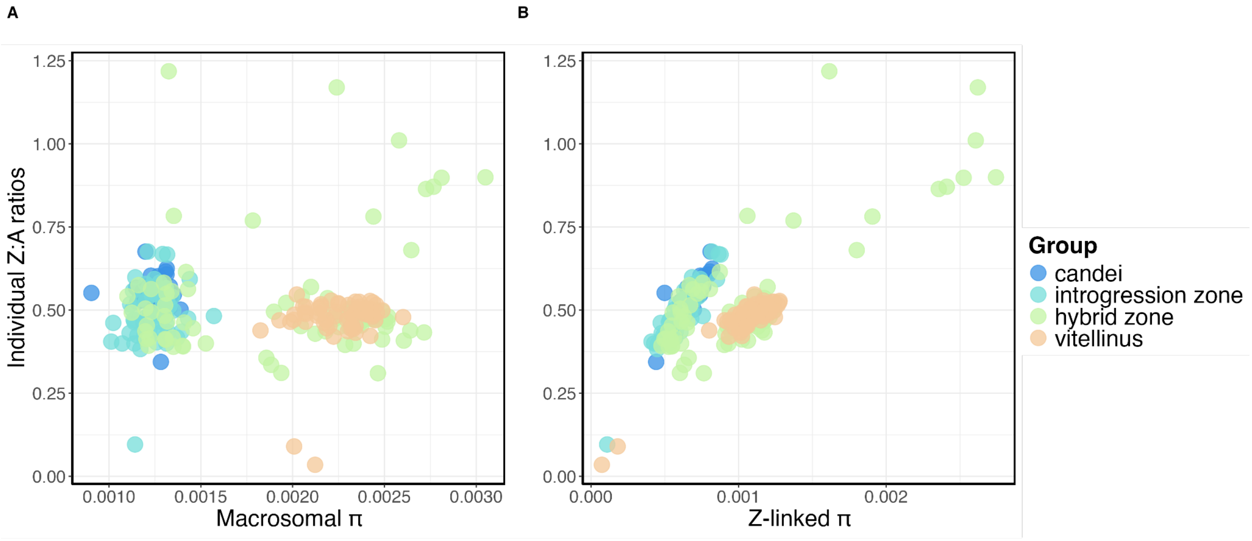


**Figure S7. High Z:A ratios in the hybrid zone are driven primarily by high Z-linked diversity. (A)** Z:A ratios vs. macrosomal π. Individuals with high Z:A ratios in the hybrid zone center (populations 8 and 9) do not, in general, have unusual levels of macrosomal nucleotide diversity. **(B)** Z:A ratios vs. Z-linked π. The individuals with high Z:A ratios in the hybrid zone center do have unusually high Z-linked nucleotide diversity.

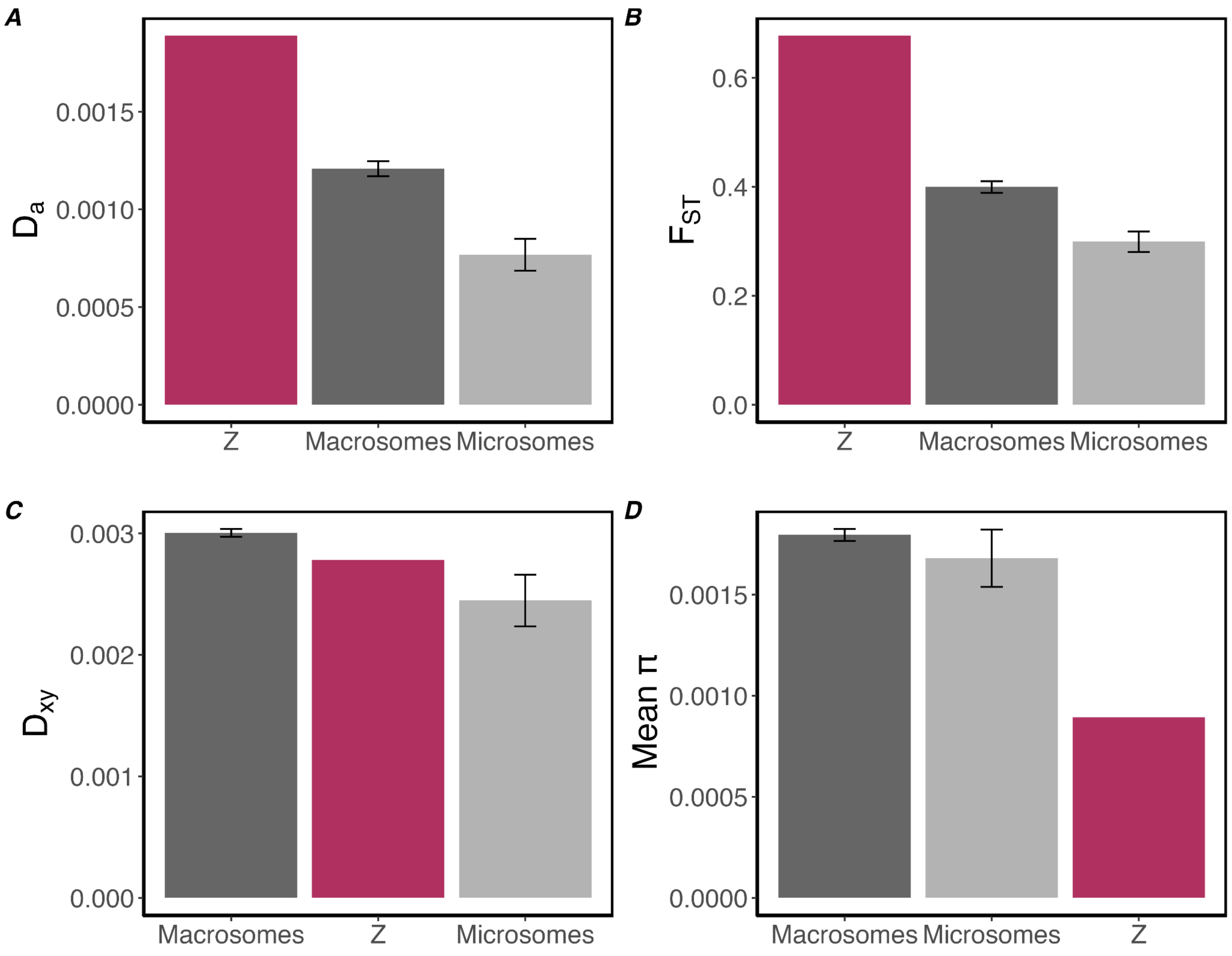


**Figure S8. Evidence for the fast-Z effect in *Manacus*.** Parental *M. candei* and *M. vitellinus* exhibited greater divergence on the Z chromosome than the autosomes based on relative measures of population differentiation. For this analysis, parental *M. candei* (populations 2 and 3) was compared to parental *M. vitellinus* (populations 9.5, 10, and 11) using *pixy* v. 2.2.1 (Korunes & Samuk, 2021). In all panels, macrosomal and microsomal bars represent the average values (± 1 SEM) across all macrosomes (n = 14) and microsomes (n = 23). Mean π is calculated by averaging the nucleotide diversity of the two parental species for each chromosome. **(A)** Net divergence ($D_a$) between parental species was greater for the Z chromosome than the two autosomal compartments. Net divergence ($D_a = D_{xy} – (\pi_1 + \pi_2)/2$) aims to estimate the amount of differentiation that has occurred since divergence from a common ancestor, which can be a more informative measure of divergence than $D_{xy}$ when lineages recently split and had significant ancestral polymorphism (Cruickshank & Hahn, 2014). **(B)** Based on Weir and Cockerham's $F_{ST}$, the two parental species are more differentiated on the Z than on the macrosomes or the microsomes. **(C)** Based on absolute divergence ($D_{xy}$), the Z shows slightly lower differentiation than the macrosomes. This is likely due to the large differences in **(D)** mean π between Z chromosomes and the autosomes. Indeed, previous work in *Manacus* has shown that $D_{xy}$ and nucleotide diversity are tightly correlated (Pearson's $r$ > 0.98; Lim et al., 2024), leading to $D_a$ being favored as a measure of divergence in this system.

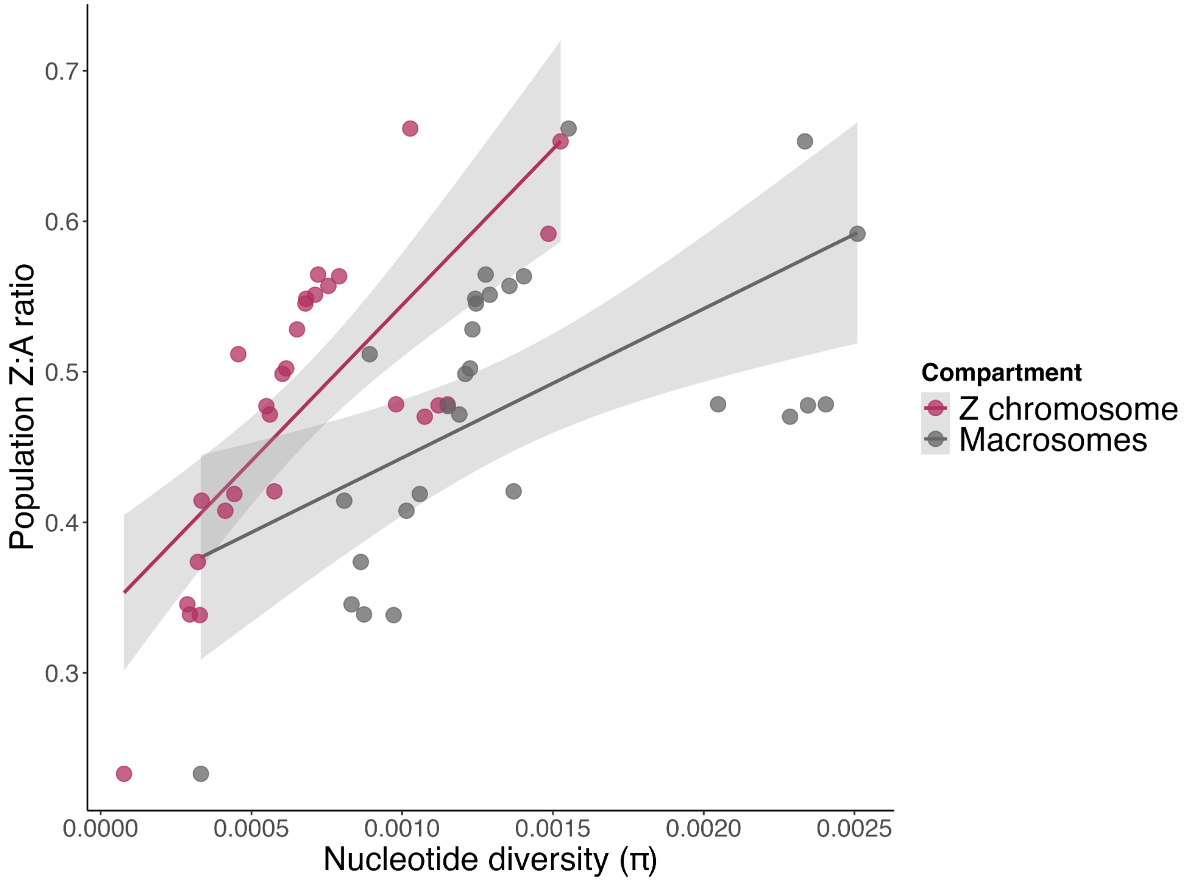


**Figure S9. Relationship between Z:A ratios and nucleotide diversity at Z-linked and macrosomal loci.** Points represent population-level π and Z:A values. Declines in overall nucleotide diversity are associated with disproportionate loss of population Z-linked diversity relative to macrosomal diversity, evidenced by the steeper slope of the Z chromosome regression line. This effect gives rise to the overall positive relationship between population nucleotide diversity and Z:A ratios shown in Figure 6.

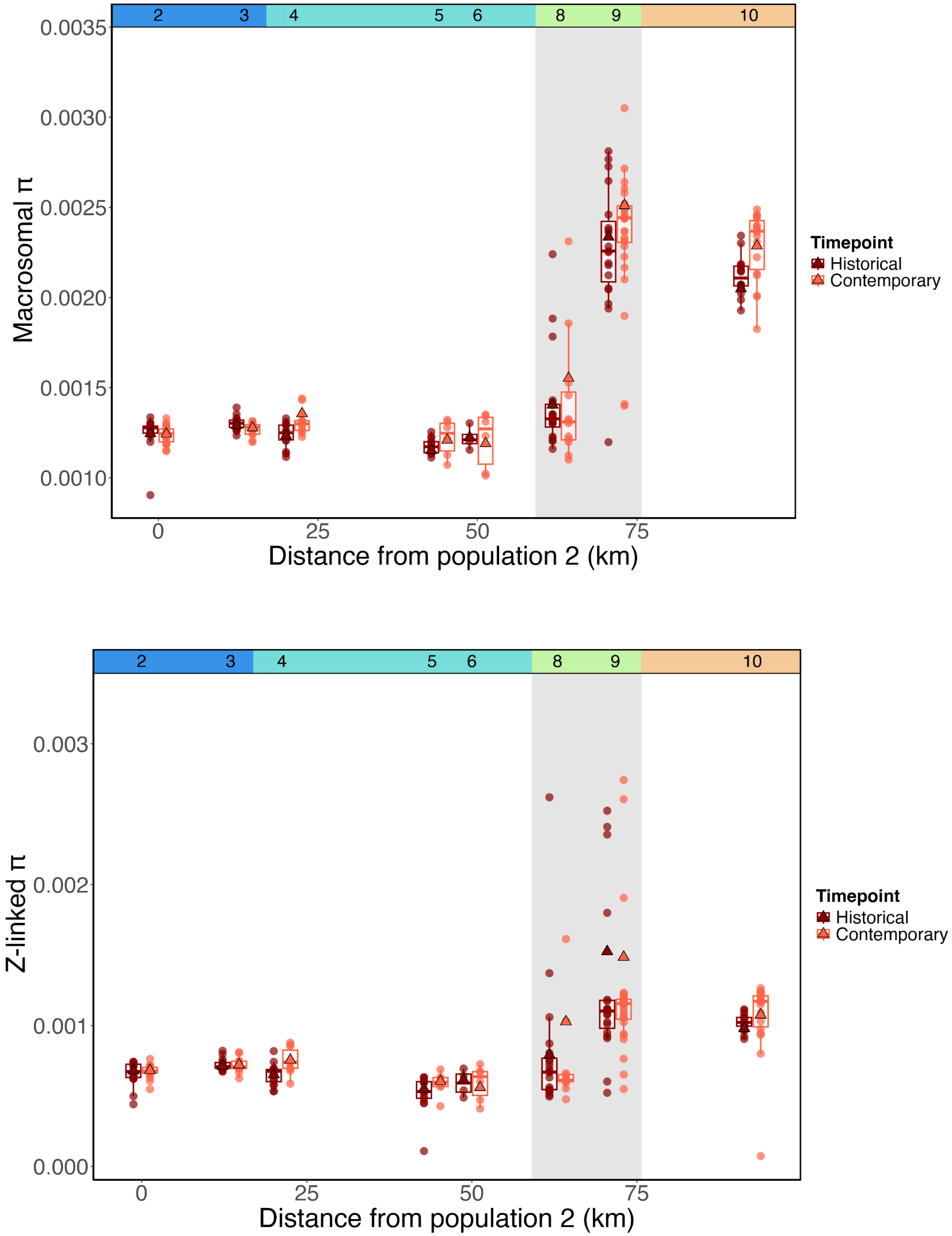


**Figure S10.** Macrosomal and Z-linked π visualized separately across the mainland transect.

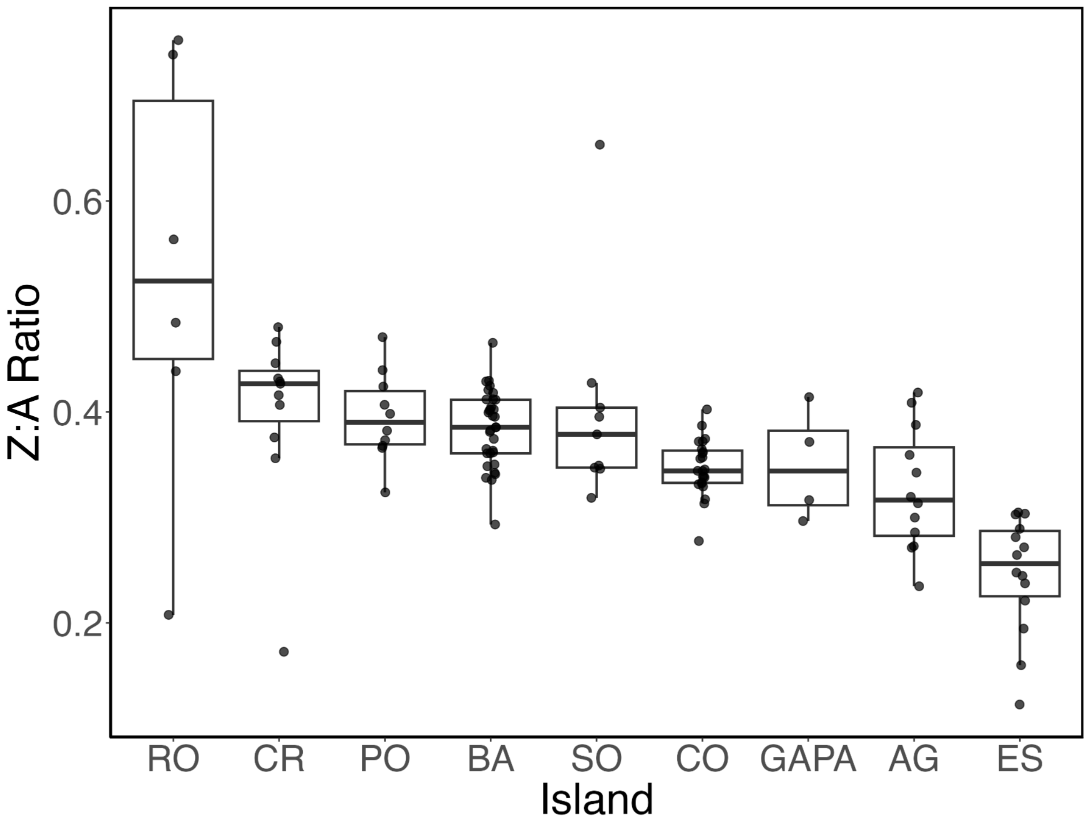


**Figure S11. Individual-level Z:A estimates for the island populations.** Island names are abbreviated as follows: Isla Cristobal (CR), Isla Pastores (PA), Isla Colón (CO), Cayo Roldan (RO), Isla Popa (PO), Isla Bastimentos (BA), Isla Solarte (SO), Cayo de Agua (AG), and Escudo de Veraguas (ES). Islands are arbitrarily ordered along the x-axis by median Z:A ratio. We note the dramatic individual-level variation exhibited in Cayo Roldan (RO), which does not fit the otherwise relatively consistent degree of within-population variation and low median Z:A estimates relative to mainland populations.

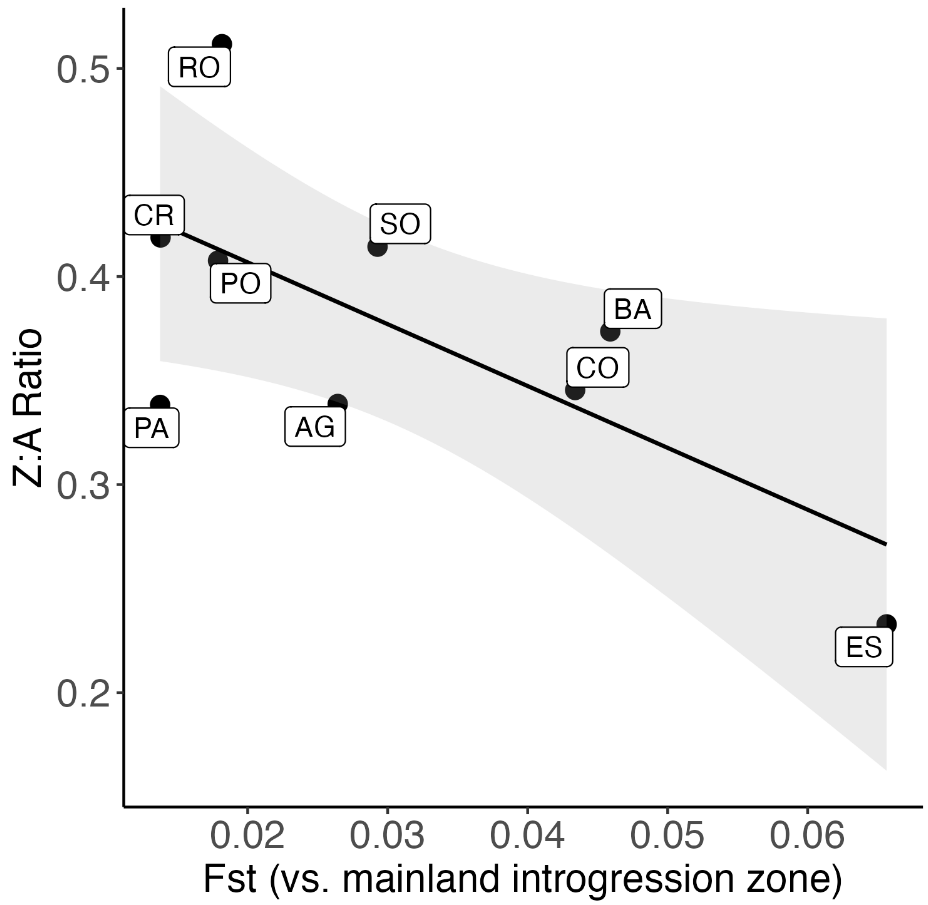


**Figure S12. Differentiation (*$F_{ST}$*) between each island and the mainland introgression zone populations.** Mean $F_{ST}$ values were obtained by comparing the population on each island to the combined populations of the mainland introgression zone (populations 4–7) in *Stacks* populations. The island populations are derived from the mainland introgression zone (see Figure 2B), and thus the introgression zone populations taken together are likely the best point of comparison for the contemporary island populations. Differentiation from introgression zone populations ($F_{ST}$) is significantly associated with the Z:A ratio (LM: t = –2.52, p = 0.04; multiple $R^2$ = 0.48), although this effect is primarily driven by the isolated island of Escudo, which has both an especially low Z:A ratio and high $F_{ST}$.

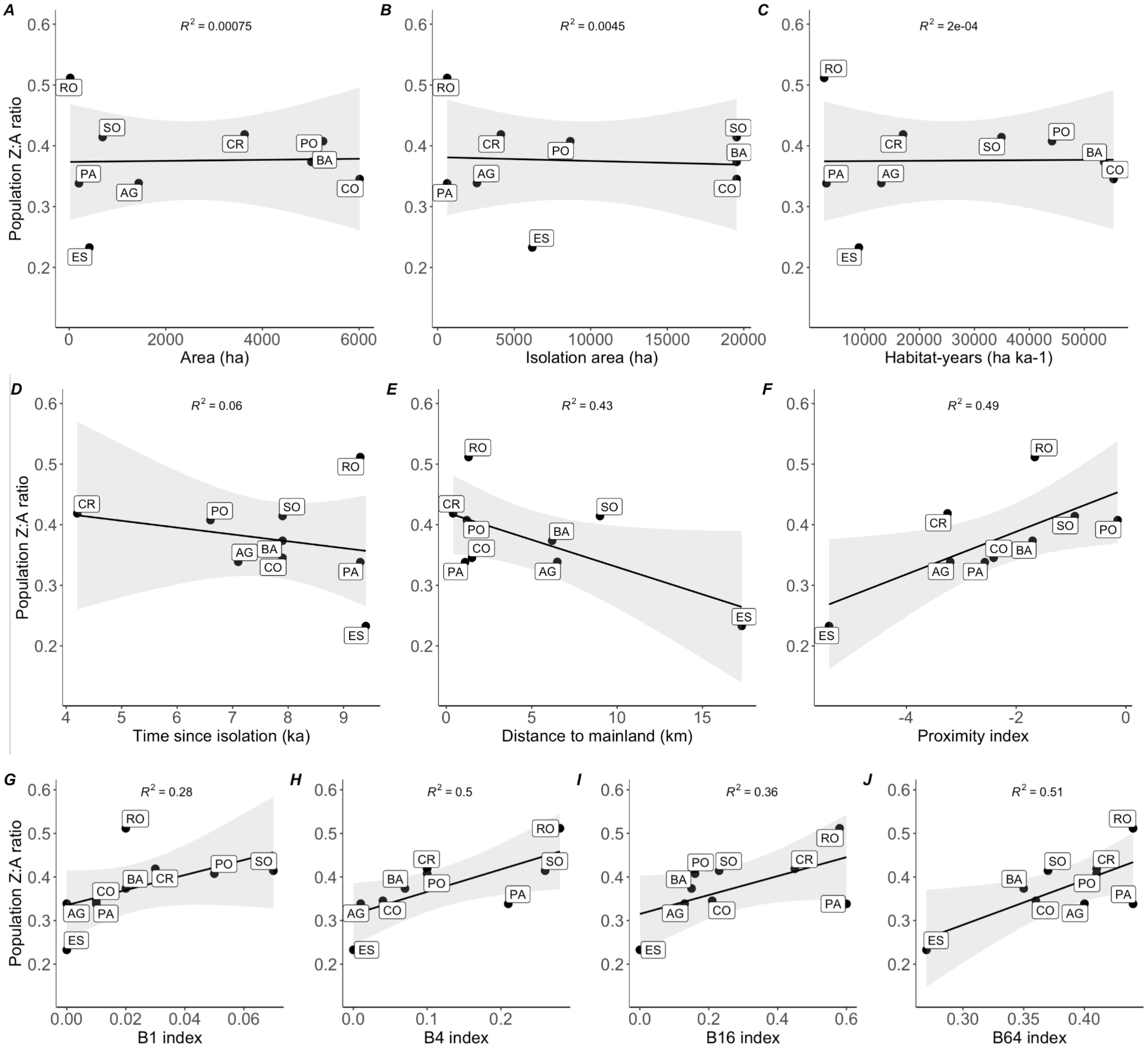


**Figure S13. Z:A ratio plotted against all individual geomorphometric variables.** Statistical outputs of univariate regression models can be found in Table S2.
